# Deciphering and steering population-level response under spatial drug heterogeneity on microhabitat structures

**DOI:** 10.1101/2025.02.13.638200

**Authors:** Zhijian Hu, Kevin Wood

## Abstract

Bacteria and cancer cells inhabit spatially heterogeneous environments, where migration shapes microhabitat structures critical for colonization and metastasis. The interplay between growth, migration, and spatial structure complicates the prediction of population responses to drug treatment—such as clearance or persistence—even under the same spatially averaged growth rate. Accurately predicting these responses is essential for designing effective treatment strategies. Here, we propose a minimal growth-migration model to study population dynamics on discrete microhabitat structures under spatial drug heterogeneity. By applying a kernel transformation, we map the original structure to an effective fully connected graph and derive a new exact criterion for population response based on a regularized Laplacian kernel reweighted by local growth rates. This criterion connects to forest closeness centrality and yields analytical bounds and sufficient conditions for population growth or decline. We find that higher structural connectivity—like increased migration—generally promotes decline. Our framework also informs optimal spatial drug assignments, which reduce to selecting interconnected subcores in the effective complete graph. For partially controllable microhabitats or unknown drug distributions, we identify strategies that ensure population decline. Overall, our results offer a new theoretical perspective on drug response in spatially structured populations and provide practical guidance for optimizing spatially explicit dosing strategies in heterogeneous environments.

## Introduction

Healthcare-related cell communities, such as bacterial infections and cancer, have increasingly been recognized as complex ecosystems over the past decades [1–3]. These systems exhibit emergent collective behaviors, including antibiotic resistance and cancer invasion, which arise from the interplay of ecological and evolutionary processes. The complex interactions between different phenotypes and genotypes have inspired novel therapeutic approaches, such as containment strategies and adaptive therapies [4–8], which leverage competition between cell types. From a dynamical systems perspective, drugs constitute an integral part of the environment, and their interactions with cells—particularly under spatially heterogeneous drug distributions—have gained attention in recent years [5, 9–39].

At the evolutionary timescale, studies on cancer have demonstrated that spatial drug heterogeneity can accelerate the evolution of drug resistance [10], shortening the time between initial treatment and the emergence of resistant clones at detectable levels. This phenomenon has been captured in minimal two-well models and extended ladder models, where differences in drug concentration modulate growth rates [16, 17]. Similar acceleration effects have been observed in bacterial populations through laboratory experiments [11, 12], though recent theoretical work using mean first passage time models suggests that spatial heterogeneity can either accelerate or decelerate resistance evolution depending on mutation and migration rates [10, 36, 39, 40]. These contrasting outcomes underscore the complexity of resistance evolution and suggest that optimal spatial drug heterogeneity design could help slow resistance development.

At the ecological timescale, similar variability is observed in patient outcomes. Even under identical drug dosages, patients exhibit different levels of pathogen or cancer cell clearance between the first treatment and the final day of therapy. For instance, despite a standard 14-day antibiotic treatment regimen, approximately 20% of patients fail to achieve sufficient clearance of H. pylori [41]. Similarly, cancer patients face recurrence risks due to insufficient clearance [5, 42, 43], which may be influenced by spatial heterogeneities in drug absorption and pharmacokinetics/pharmacodynamics (PK/PD) [25]. Unlike well-mixed laboratory conditions, the in vivo tumor and microbial environments exhibit spatial complexity, making traditional well-mixed drug-dose response measurements insufficient for predicting clinical outcomes. Recent work [44] has modeled population-level responses as an eigenvalue problem, where cell survival or extinction is governed by the interplay of spatial growth, migration, and microhabitat connectivity in a one-dimensional setting. This framework has been extended to model the emergence of multidrug resistance under natural selection [45]. However, the entanglement of structural, migratory, and growth-related factors in these eigenvalue formulations limits their interpretability. Additionally, clinical applications require extending these results beyond one-dimensional models to account for multiple discrete microhabitat scenarios such as cancer metastasis [46–51], lymph node networks [52–54], and bacterial translocation across body compartments [15, 55, 56].

To better understand how spatial drug heterogeneity shapes population-level responses—specifically, whether cell populations grow or decline—we develop a minimal growth-migration model defined on discrete microhabitat networks. In this model, local growth rates are constrained by drug-induced limits, ranging from a maximum drug-free growth rate to a minimum death rate. Migration allows for population movement across interconnected microhabitats, embedding spatial structure into the dynamics. We theoretically analyze this model by transforming the original network into an effective fully connected graph using a regularized Laplacian kernel. This transformation enables the derivation of an exact criterion for determining population response, reweighted by local growth rates and interpreted through a centrality measure known as forest closeness centrality. This centrality interpretation provides an efficient and analytically tractable condition for predicting whether a population will persist or be cleared under a given spatial drug distribution.

Interestingly, we find that increasing global structural diffusion, such as through enhanced migration or more inter-microhabitat connectivities, leads to a smooth transition from population growth to decline when the spatial drug heterogeneity is fixed. For a given microhabitat structure, population dynamics are not only governed by the spatially averaged growth rate and the migration rate, but the spatial arrangement of growth rates still plays a critical role. Our framework informs optimal spatial drug assignments, which reduce to selecting interconnected subcores in the effective complete graph. For partially controllable microhabitats, we can design optimized strategies for population clearance; for unknown spatial drug heterogeneities, under a centrality-based heuristic, we identify parameters that ensure robust population decline, independent of spatial drug arrangements.

These findings have important implications for understanding and manipulating drug responses in complex biological systems. Health-related applications—such as bacterial infections and metastatic cancer—often involve heterogeneous spatial environments where growth, migration, and structure interact in nontrivial ways. Our framework offers a generalizable and analytically grounded approach to assess when populations will decline in such settings, providing a new lens through which to interpret treatment outcomes. Moreover, the model serves as a foundation for future extensions, including those incorporating temporal fluctuations, environmental feedback, or interspecies interactions. Clinically, this work highlights the potential of spatially explicit dosing strategies, tailored not only to drug strength but also to the structure of the tissue or infection site, to improve treatment outcomes for both infections and solid tumors.

### Basic set-up of model system

Cell populations like bacteria, or cancer cells, can actively or passively move between different sites by migration or metastasis. When accounting for fluctuations in population growth and the stochastic nature of drug-induced clearance [57], the system can be described using stochastic microscopic dynamics governed by a master equation (see SI). However, in human bodies, where bacterial and cancer cell populations typically exceed thousands, we can apply the Van Kampen approximation [58]. This allows us to ignore microscopic dynamic details and derive a simplified macroscopic growth-migration equation

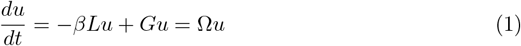

This is equivalent to a linearized exponential growth from logistic growth, by ignoring the competition or density-dependent effect, where

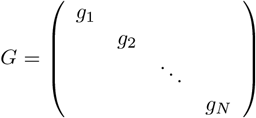

is a diagonal matrix describing growth rates induced by drug at different microhabitats. And *L* is the Laplacian matrix describing the multi-microhabitat structure. *β* describes the migration rate and *u* = (*u*_1_, *u*_2_, …, *u*_*N*_)^*T*^ describes the population densities on different microhabitats. *N* is the total number of microhabitats. Since drug-induced growth rates {*g*_*i*_} cannot be infinite, we require *g*_*i*_ ∈ [*d*_0_, *g*_0_], where *d*_0_ < 0 is the maximum death rate, and *g*_0_ > 0 is the maximum growth rate or drug-free growth rate, depending on drug type and cell type, or other environmental factors like nutrition [59–61]. For example, single bactericidal drug like Ampicillin can be fitted into a hill fuction to get maximum growth rate at a drug-free environment, or the maximum death rate when the AMP concentration is extremely high [62]. Equation (1) is the minimal growth-migration model required to understand short-term population response with different spatial drug heterogeneities and different multi-microhabitat strcutures. Since the drug concentration must be sufficiently high to eliminate cancer or bacterial cells, we require that the spatially averaged growth rate satisfies 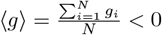. For population decline or clearance, the condition Ω = *G* − *βL* ≺ 0 must hold, or equivalently, the largest eigenvalue *λ*_0_ = ∥Ω∥ = ∥*G* − *βL*∥ < 0. However, solving this exactly is typically intractable. Applying first-order perturbation theory, as in recent studies [44, 45], we approximate the largest eigenvalue as

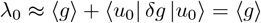

Since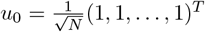, we obtain ⟨*u*_0_| *δ*_*g*_ |*u*_0_⟩ = 0, indicating that this approximation only captures well-mixed effects while neglecting spatial drug distribution and multi-microhabitat structure. Higher-order approximations may be necessary but are challenging to compute.

## Results

### Kernel transformation and new criterion for population response

Instead of relying on perturbation theory, we derive an exact solution for the clearance or decline criterion based on a “kernel” transformation. For the kernel transformation, we first decompose *G* as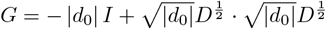, where *I* is the identity matrix and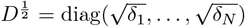is an orthogonal diagonal matrix with the ith diagonal entry as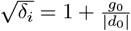, representing the relative growth information over microhabitats compared to the maximum death rate |*d*_0_|. Based on Schur complements (see SI for details), we show that the condition Ω = *G* − *βL* ≺ 0 is equivalent to

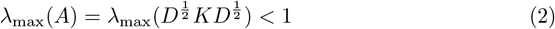

where *K* = (*I* + *γL*)^−1^ is the regularized Laplacian kernel matrix naturally emerging from our minimal growth-migration dynamics, and 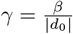 is the ratio of migration rate to maximum death rate. Traditionally, the parameter *γ* characterizes the extent of diffusion or spread over the graph structure. Here, it effectively acts as a rescaled migration intensity, modulating how spatial structure shapes population outcomes.

The matrix *K*, also known in some contexts as the *forest matrix* or the *parametrized matrix forest index*, has appeared in a wide range of network applications. It is at least row-stochastic (and becomes doubly stochastic for undirected graphs), mapping the original sparse structure into an effective, fully connected graph. Off-diagonal entries *K*_*ij*_ have been used to quantify similarity between spatial locations in disease-spread models, such as COVID-19 transmission networks [63]. More broadly, *K* has been applied for link prediction in diverse domains, including social and biological networks [64], protein–protein interaction (PPI) networks [65, 66], and virus–host interaction inference [67]. A particularly relevant measure derived from *K* is the *forest closeness centrality* [68, 69], *defined as*

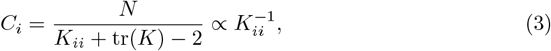

which approximately corresponds to the inverse of the diagonal element *K*_*ii*_. For simplicity, we will refer to 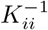 as the centrality measure in this paper. While this matrix has traditionally been engineered for tasks such as node ranking, link prediction, or similarity assessment, our framework reveals its *natural appearance* as a kernel governing the interplay between growth and migration. This connection provides a simple but novel interpretation of centrality as a predictor of population response and optimized clearance strategies, which we further illuminate in the subsequent sections.

Rewrite 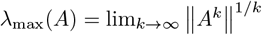 where ∥ · ∥ is chosen as the infinite norm | · |_∞_, the maximum row sum of absolute value of matrix *A*. 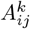 here can be interpreted as the length-k walk from microhabitat *i* to microhabitat *j*. So now *λ*_max_(*A*) becomes a special norm measuring the maximum total walk length from a microhabitat *i* to every microhabitat including *i* itself (see SI for details). Again, this tells us the new largest eigenvalue is capturing global information of the original multi-microhabitat structure. It’s hard to find out the analytical solution of *λ*_max_(*A*) due to the inhomgeneous nature of growth-weight distance between microhabitats. However, based on this walk-length interpretation, we find out the lower and upper bound for *λ*_max_(*A*),

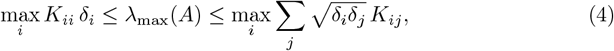

where *K*_*ii*_ ∝ 1*/C*_*i*_ in the lower bound represents the inverse of forest closeness centrality, and the upper bound is the maximum row sum of length-1 walk (see SI for details). We can also treat each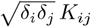 as the edge value of the efficient fully connected graph.Then these 2 bounds inform that, finding *λ*_max_(*A*) here is approximately equivalent to identifying key microhabitats with maximum growth-weighted centrality, or with maximum weighted connectivities. *λ*_max_(*A*) = max_*i*_ *K*_*ii*_ *δ*_*i*_ holds when microhabitat *i* becomes effectively isolated;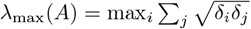 holds when the structure has local symmetry and *A* becomes a circulant matrix(see SI for details). Either *K*_*ii*_ or *K*_*ij*_ contains global information of the original structure due to the kernel transformation. If we do the same calculation for Ω = *G* − *βL* instead of *A*, we can only find out *g*_*i*_ − *βk*(*i*) ≤ *λ*_max_(Ω) ≤*g*_*i*_, where *k*(*i*) is the degree of ith mcriohabitat. These are looser bounds only containing localized information like degree. So by doing the kernel formation, we can find out a tighter bound of *λ*_max_ with meaningful interpretations.

If 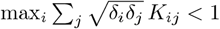 or max_*i*_ *K*_*ii*_ *δ*_*i*_ > 1, we thus find out sufficient conditions for population decline or population growth (or 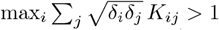, max_*i*_ *K*_*ii*_ *δ*_*i*_ < 1 are necessary conditions for population growth, population decline). These 2 criteria can be applied to quickly determine population response without directly calculating *λ*_max_(*A*) itself. While 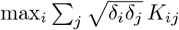 is in a first-order form considering interaction effect, max_*i*_ *K*_*ii*_ *δ*_*i*_ has zero-order form considering only centrality effect, decoupling growth information and structure information^1^.

So in practice, we can always perform a quick diagnostic check of the population response using the quantity max_*i*_ *K*_*ii*_ *δ*_*i*_. If max_*i*_ *K*_*ii*_ *δ*_*i*_ > 1, we can immediately predict population growth. Conversely, if max_*i*_ *K*_*ii*_ *δ*_*i*_ < 1, this provides a necessary condition for population decline. As a second step, we evaluate the upper bound 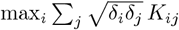; if this quantity is also smaller than 1, we can conclude that population decline will occur. Since the diagonal element *K*_*ii*_, also known as the forest closeness centrality *C*_*i*_, serves as a useful and interpretable indicator under growth-migration dynamics, we propose to refer to *K*_*ii*_ as the *inverse dynamic-related centrality*, and denote its reciprocal *C*_*i*_ as the *dynamic-related centrality*.

In this study, we primarily focus on how inverse centrality values *K*_*ii*_ can be leveraged to predict population responses. Because these responses are influenced by two spatial effects — spatial drug heterogeneity (i.e., the spatial growth distribution {*δ*_*i*_}) and the spatial structure of the system — we organize our analysis as follows:

- Given a fixed spatial drug heterogeneity {*δ*_*i*_}, we analyze how different microhabitat structures, as captured by the matrix *K*, influence population-level responses, and examine the role that *K*_*ii*_ plays in shaping these dynamics.
- Given a known multi-microhabitat structure (i.e., a fixed matrix *K*), we explore how different growth distributions {*δ*_*i*_} affect the outcome, and how this insight can be used to optimize treatment strategies by exploiting structural information — particularly the inverse centralities *K*_*ii*_.

### Microhabitat structure effects under given spatial drug heterogeneity

We aim to investigate how microhabitat structures influence population responses under a fixed spatial drug heterogeneity. Specifically, we address the following questions: 1. How do different microhabitat structures with the same connectivity affect population responses? 2. How does varying the level of connectivity alter spatial drug responses across different structural configurations? Overall, our goal is to understand how structural effects on population dynamics can be predicted by the dynamic-related centrality *C*_*i*_, or equivalently, by its inverse *K*_*ii*_.

### Distinct population responses induced by different multi-microhabitat structures predicted by kernel criteria

Starting from a fixed spatial drug heterogeneity (see Figure 2A), we demonstrate how different multi-microhabitat structures lead to distinct population responses, which can be precisely captured by checking max_*i*_ *K*_*ii*_ *δ*_*i*_ and 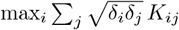. The inverse value of dynamic-related centralities *C* are visualized for two different multi-microhabitat structures, with color gradients indicating different values (see

Figure 1B). All values of inversed *C* (or *K*_*ii*_) are upper bounded by 1 (see SI). Different transparencies represent varying growth rate values.

**Figure 1.**
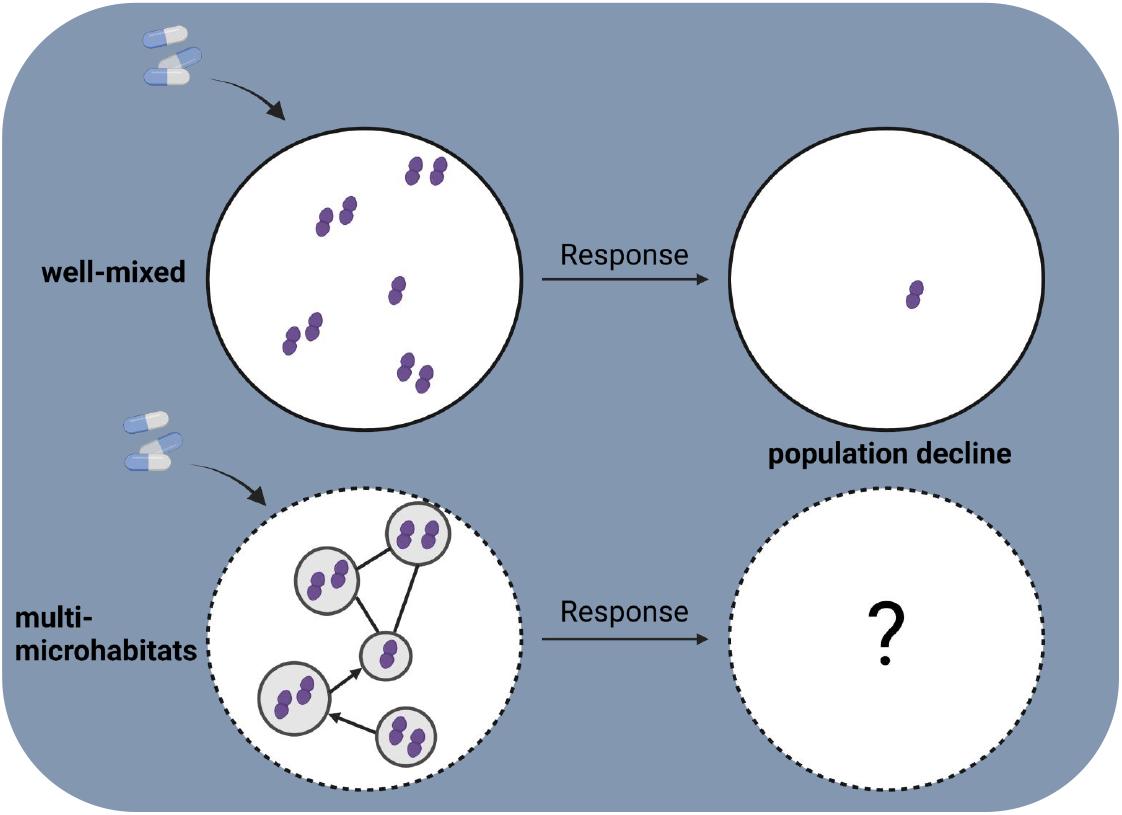
Illustration of Population Response Under Drug Exposure. Example of multiple bacterial microhabitats and their diverging population responses under high bactericidal drug doses. Despite receiving the same drug dose, bacterial populations may exhibit microhabitat-dependent responses, even when the population would decline in a well-mixed environment.

**Figure 2.**
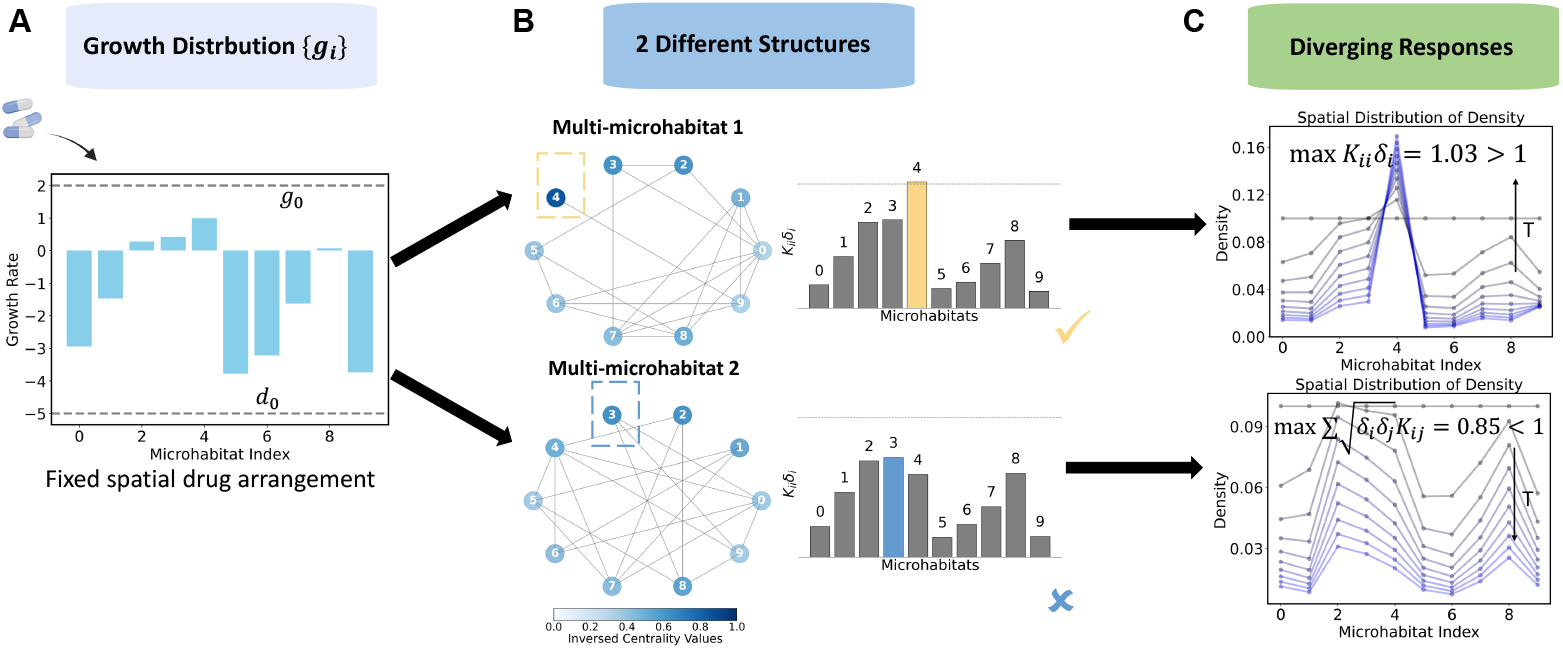
Different microhabitat structures with the same spatial growth distributions induce different population responses, explained by our criterion. **A**. Example distribution of spatial growth rates across 10 connected microhabitats. Growth rates are bounded by the drug-related maximum growth rate and death rate, represented by grey dashed lines. **B**. Two different microhabitat structures with the same number of microhabitats and migration connections, along with their corresponding *K*_*ii*_*δ*_*i*_ values shown as bar plots. In each structure, colors indicate *K*_*ii*_, the inverse of dynamic-related centrality values *C*_*i*_, while different transparencies represent relative growth *δ*_*i*_. The microhabitat with the highest *K*_*ii*_*δ*_*i*_ in each structure is highlighted with a triangle, colored light yellow for population growth and light blue for population decline, as confirmed by the adjacent bar plots. Notably, the microhabitat with the highest *K*_*ii*_ does not necessarily correspond to the highest *K*_*ii*_*δ*_*i*_. The yellow check mark near the top bar plot is showing that the population growth can be predicted by checking max_*i*_ *K*_*ii*_*δ*_*i*_ > 1. The light blue cross mark near the bottom bar plot indicates that max_*i*_ *K*_*ii*_*δ*_*i*_ < 1 serves as a necessary condition for population declinew, and further check of 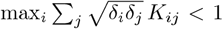 is required. **C**. Two spatial dynamic configurations demonstrating different population responses. The grey dashed line represents the initial population density. As time increases, the density curves change from grey to dark blue. The dynamics induced by structure 1 exhibit a growth trend, consistent with our theoretical prediction, while those induced by structure 2 show a decline trend predicted by 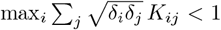. For both 2 structures, *β* = 1, *g*_0_ = 2, *d*_0_ = −5, ⟨*g*⟩ = −1.5, and the edge number is fixed at |*E*| = 10.

Notably, the locations of *g*_max_ (or *δ*_max_), max *K*_*ii*_, and max_*i*_ *K*_*ii*_ *δ*_*i*_ do not necessarily coincide. We can approximate *K*_*ii*_ *δ*_*i*_ directly from the visualized networks by observing the color and transparency of microhabitats(nodes), which are highlighted in different colors. The adjacent bar plots reveal that in one structure, at least one *K*_*ii*_ *δ*_*i*_ value exceeds 1, whereas in the other, all values remain below 1. Consequently, the first multi-microhabitat structure leads to population growth, while the second indicatively results in decline. It’s proved by checking 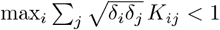. The population decline is as illustrated by the spatial-temporal dynamics in Figure 2C. Our simple criterion thus serves as a powerful predictive tool for population responses without requiring detailed dynamic simulations. Interestingly, dynamic-related centrality *C* itself, is proportional to node degree, correlating with many conventional centrality measures (see SI), although the importance rankings are not exactly the same. It bears strong similarity to closeness centrality. In the population responses of 2 example structures we haven shown, structure 2 with more even connectivities for different microhabitats, tend to decline compared to structure 1. This can be explained from a centrality perspective: high *C* usually represents good connectivities with other microhabitats, if under a relatively high-drug dose regime as shown in Figure 2A, the high *C* tends to connect with more “sink” microhabitats with negative growth rates, increasing the probability for population decline. In structure 1, the microhabitat 4, with positive growth rate high inversed centrality value (low centrality *C* value), is pretty isolated with only one connection to other microhabitats, thus increasing the chance of survival.

### Increased connectivity accelerates centrality-predicted population decline

Since dynamic-related centrality reflects the importance of edge number, and the second structure with a more even connectivity in Figure 2 leads to population decline, it suggests that increasing global diffusion over different microhabitats can effectively decrease *K*_*ii*_, thus mitigating population growth by max *K*_*ii*_*δ*_*i*_ < 1. Increasing migration rate can be one way to enhance diffusion(we can also see that from the mathmatical form of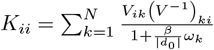, whee *K*_*ii*_ ∝ *β*^−1^). Increasing the average number of edges or connection density may be another way to enhance global diffusion thu increasing the probability of population decline. To investigate this, we tune the edge density 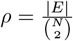, defined as the ratio of the total number of edges in a given graph to the number of edges in a complete graph, and examine its effect on max *K*_*ii*_*δ*_*i*_.

Using 1000 stochastic samples from random graph for each edge density, we observe that as edge density increases, sample-averaged max *K*_*ii*_*δ*_*i*_ decreases rapidly and eventually crosses the critical threshold max *K*_*ii*_*δ*_*i*_ = 1 (see Figure 3). Figure 3A illustrates four different multi-microhabitat structures with increasing edge densities, where the colors gradually fade to white, indicating decreasing max *K*_*ii*_*δ*_*i*_ values. This finding suggests that densely connected microhabitats may require lower drug doses to induce population decline, whereas sparsely connected structures may necessitate higher drug doses to achieve the same effect.

**Figure 3.**
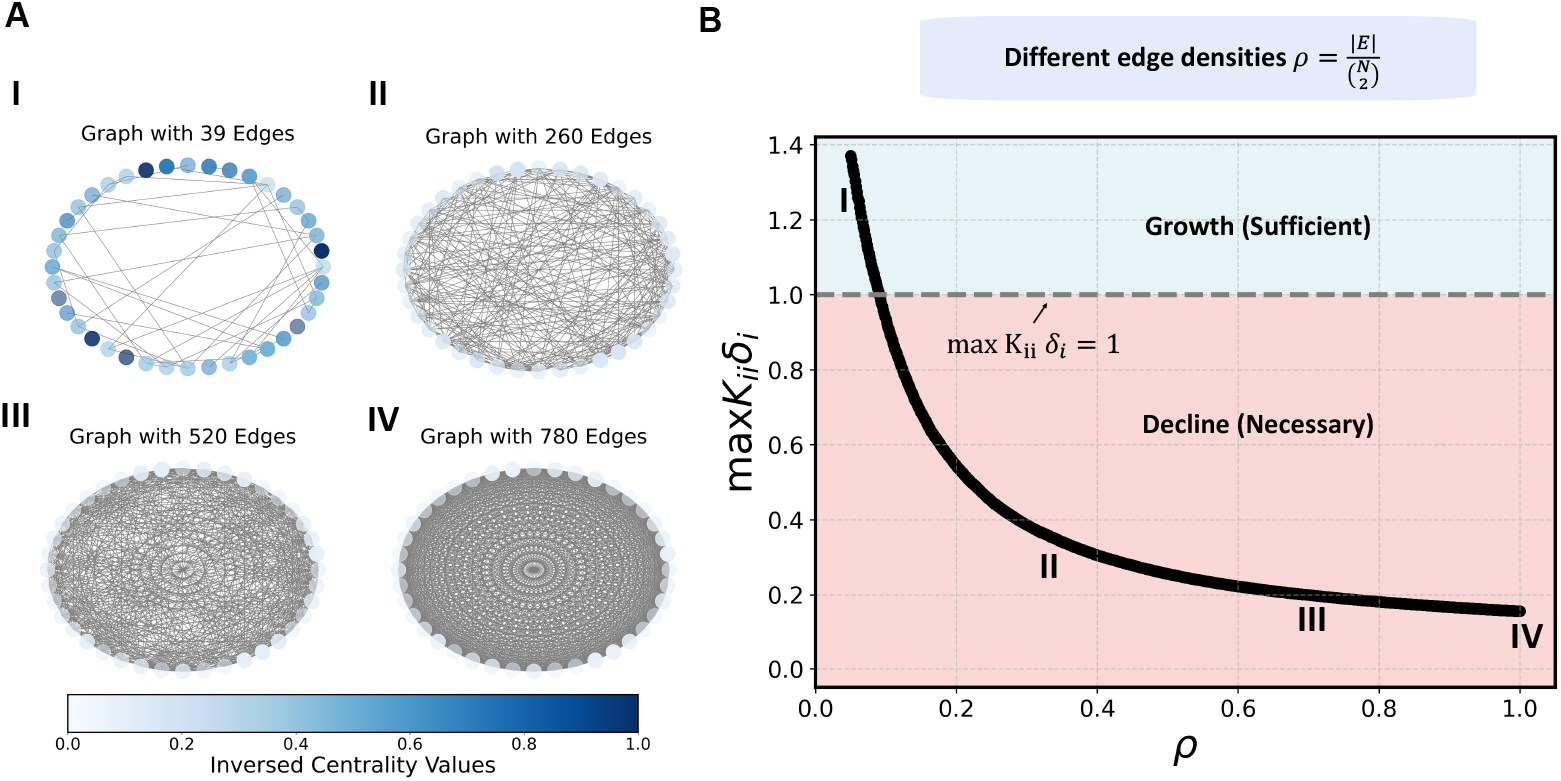
Increasing edge/connection density enhances the tendency for population decline. A. Four example microhabitat structures(microhabitat number *N* = 40) with varying connection densities but the same spatial growth rate distribution. Color gradients represent dynamic-related centrality values *C*, while different transparencies indicate relative growth values *δ*. As connectivity increases, the colors gradually fade to white, indicating lower centrality values. **B**. The maximum *Cδ* value decreases as connection density *ρ* increases. As the curve crosses the critical threshold max *Cδ* = 1, population response shifts from growth to decline. Connectivity has a similar effect to migration rate (see SI). The curve represents an average over 1000 stochastic samples of different random networks for each edge density. For parameters, *β* = 0.5, *g*_0_ = 2, *d*_0_ = −1, ⟨*g*⟩ = −0.25.

### Spatial drug heterogeneity effects and optimal strategy under a given multi-microhabitat structure

Different spatial drug heterogeneities, like different multi-microhabitat structures, induce distinct population responses. In our study, to allow meaningful comparison across scenarios, we control the spatially averaged drug dose - or equivalently, the spatially averaged growth rate ⟨*g*⟩ - in our minimal model. We consider two cases depending on whether spatial drug heterogeneity is controllable: 1. Fully controllable heterogeneity: Clinically, if we are fortunate enough to precisely control drug concentration at each microhabitat (or node), population decline can be achieved by selecting the spatial drug assignment that minimizes the largest eigenvalue - we simply find when min *λ*_max_(*A*) < 1. This gives us the optimal spatial drug configuration among all possible combinations. 2. Uncontrollable or unknown heterogeneity: If spatial drug assignment cannot be precisely controlled or is unknown, we must ensure population decline under the worst-case spatial drug configuration. This requires max *λ*_max_(*A*) < 1, ensuring a robust population decline that is independent of the specific drug distribution.

This naturally leads to a constrained optimization problem, as illustrated in recent work on robust decline under spatial drug heterogeneity in continuous 1D space [44]. In both cases, we are solving a constrained nonlinear optimization problem involving the largest eigenvalue. However, the search space is large, since each *g*_*i*_ varies continuously and the configuration space grows combinatorially with the number of microhabitats *N*.

Remarkably, the solution to this class of problems can be simplified. For the minimization case (min *λ*_max_(*A*)), assuming ⟨*g*⟩ < 0, the optimal strategy is alway an interior point with every 0 *< δ*_*i*_ *< c*, or *d*_0_ *< g*_*i*_ *< g*_0_. This optimal strategy depends on *K* and is usually hard to derive analytically. However, a uniform drug assignment: *g*_*i*_ = ⟨*g*⟩ for all microhabitats, as the “even-spread strategy”, is good enough and can always induce population decline under ⟨*g*⟩ < 0. For the maximization case (max *λ*_max_(*A*)), the optimum typically lies on or near the active constraints (see SI for a detailed proof). This means that, for a fixed ⟨*g*⟩, we should assign as many microhabitats as possible to the maximum death rate *d*_0_ or the maximum growth rate *g*_0_, leaving at most one remainder *g*_*r*_ ∈ [*d*_0_, *g*_0_]. The optimal configuration then takes the form {*g*_*i*_}_optimal_ = {*g*_*r*_; *g*_0_…, …; *d*_0_,}, which we refer to as the “one-remainder strategy”. Hence, this continuous eigenvalue optimization problem reduces to a discrete combinatorial optimization. Interestingly, our kernel transformation is able to further simplify the problem. By expressing growth in terms of relative rates *δ*_*i*_, the optimal growth configuration becomes {*δ*_*i*_} _optimal_ = {*δ*_*r*_; *c*,…; 0,…}, where *δ*_*r*_ ∈[0, *c*], and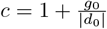 is the maximum possible relative growth rate. For nodes assigned *g*_*i*_ = *d*_0_, *δ*_*i*_ = 0, which effectively removes all edges from microhabitat *i* to its neighbors under our transformation (as the structure becomes effectively fully connected). If *N* − *S* microhabitats are suppressed (*δ*_*i*_ = 0), the remaining effective graph forms an *S*-clique, and the matrix *A* reduces to a smaller principal submatrix *A*_*S*_ of size *S*. This matrix can be decomposed as:

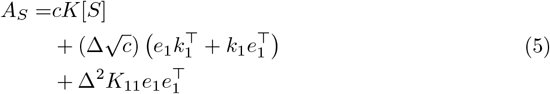

Here, we assume the microhabitat with the remainder *g*_*r*_ is labeled 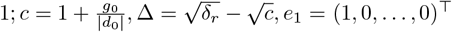, and *k*_1_ = (*K*_11_, *K*_21_, …, *K*_*S*1_)^T^ is the first column of *K*[*S*]. *K*_11_ reflects the inverse dynamic-related centrality of the node with the remainder. Thus, solving the optimization problem becomes equivalent to selecting an *S*-clique from the full graph to minimize or maximize *λ*_max_ (*A*_*S*_).

While discrete combinatorics optimization remains computationally challenging, we propose a centrality-based heuristic that leverages the decoupling of structure and growth. Under very high drug doses - i.e., when 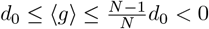, or when *g*_0_ ≥ (*N* − 1) |*d*_0_| - the one-remainder strategy becomes equivalent to selecting a single growing microhabitat. Then we have *λ*_max_(*A*) = *δ*_*r*_*K*_*ii*_. To guarantee decline under unknown heterogeneity, we require 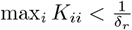, implying the structure must contain at least one relatively isolated microhabitat.

In the general case with multiple positive growth rates {*δ*_*r*_; *c*,…}, we aim to identify an interconnected but relatively isolated *S*-clique to ensure robust decline. This intuition aligns with Figure 3B, where less connected graphs tend to suppress growth less effectively, thus serving as the “worst” case. Note that selecting the most isolated S-clique is not simply equivalent to minimizing the row sum of *A*_*S*_, which is only an upper bound: 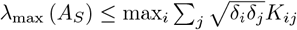.

### Optimal drug assignment with incomplete controllable microhabitats

Under complete control, the good enough strategy to minimize *λ*_max_(*A*) is to assign a uniform drug distribution: *g*_*i*_ = ⟨*g*⟩ across all microhabitats. However, in practice, we may only be able to control a subset of microhabitats and apply high enough drug concentrations to push them to the maximum death rate *d*_0_, while the remaining microhabitats stay approximately drug-free at growth rate *g*_0_. In this scenario, the problem again becomes a discrete combinatorial optimization - selecting the optimal subcore.

When the number of controllable microhabitats is large, we should avoid targeting the well-connected core. Instead, we preserve this core (i.e., assign it positive growth rates), since it can effectively “sense” the largest number of the surrounding “sink” microhabitats with maximum death rate *d*_0_. Interestingly, when the number of controllable microhabitats is very limited-e.g., we can apply high drug concentration to only one node (setting *δ*_*i*_ = 0)-we find that the optimal strategy is to suppress the microhabitat with the smallest *K*_*ii*_, i.e., the node with the highest dynamic-related centrality (which often corresponds to highest degree). With more available microhabitats to control, we continue suppressing other nodes with high centralities in descending order.

Table 1 summarizes optimized strategies across different graph families based on this principle. For more complex structures, this centrality-based rule often provides a greedy but effective approximation of the optimal solution. However, to find the true optimal drug configuration, one still needs to solve the full discrete combinatorial problem of minimizing *λ* (*A*_*S*_) with partial control.

**Table 1.**
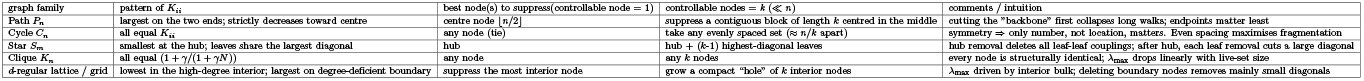
Optimized clearance strategies with limited controllable microhabitats, across different graph families based on the regularized Laplacian kernel matrix *K*.

This drug assignment strategy under limited controllability closely mirrors strategies used in epidemic outbreak suppression in susceptible-infectious-susceptible (SIS) network dynamics [70–73], where the optimal curing rate of each node is made proportional to its degree. Thus, in our minimal growth-migration model: a. when many microhabitats are controllable, we protect the well-connected core; b. when only a few are controllable, we target the core to suppress it most effectively.

### Robust population decline with unknown spatial drug heterogeneity

In most real-world scenarios, especially within the human body, estimating or controlling drug concentrations at each microhabitat is infeasible. Spatial drug heterogeneity is typically unknown (Figure 4A). Assuming we can estimate the migration rate *β* and control the total drug dose (or the spatially averaged growth rate ⟨*g*⟩), the problem becomes identifying robust clearance strategies that ensure population decline in the worst case. That is, we need:

**Figure 4.**
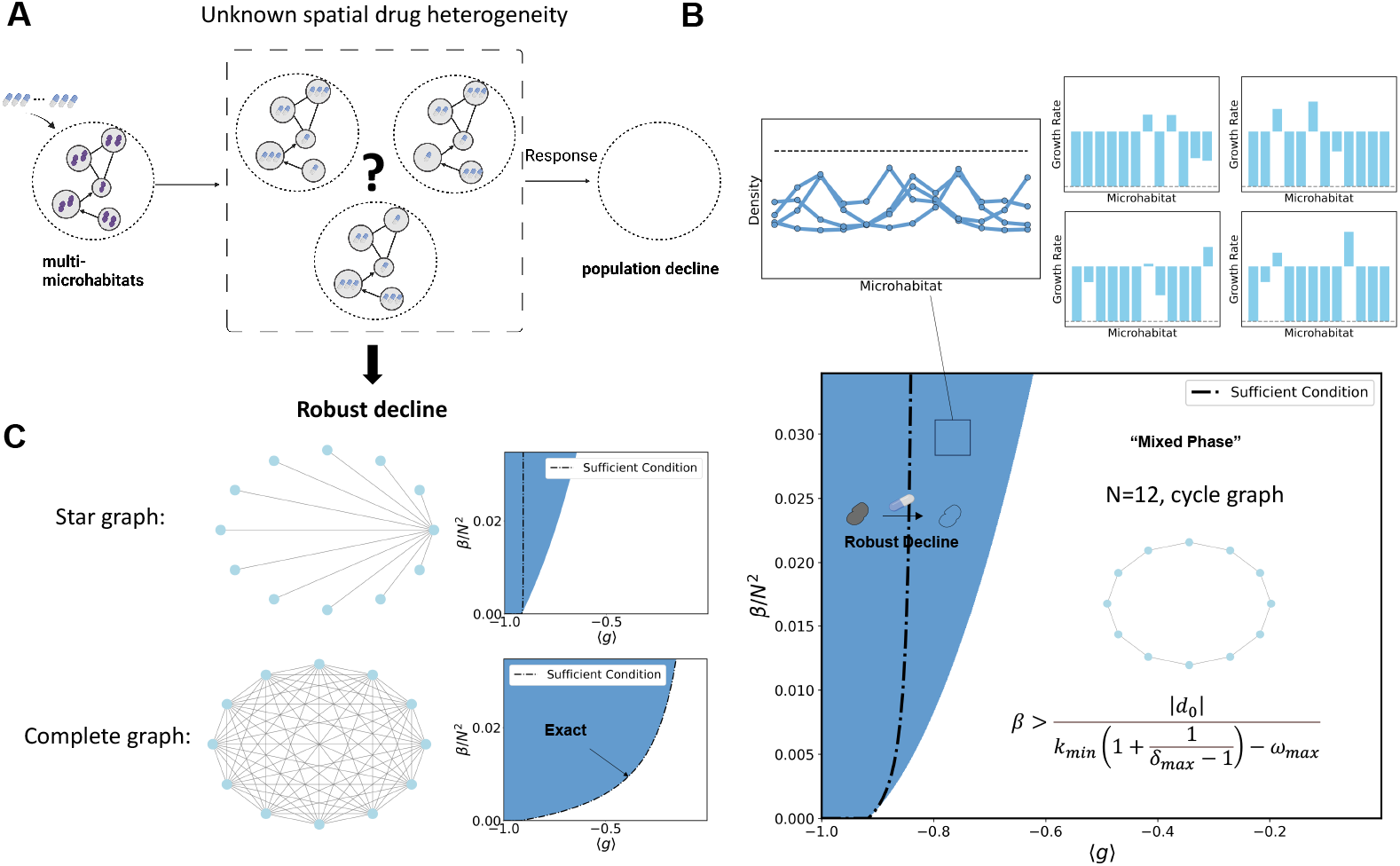
Robust population decline under unknown spatial drug heterogeneity. A. Illustration of the concept of robust population decline. When spatial drug heterogeneity is unknown, the clearance strategy evaluates whether population decline always occurs under a given migration rate and total drug dose, effectively eliminating the variability induced by different spatial drug configurations. **B**. Example phase diagram on a cycle graph, showing the relationship between migration rate *β* and spatially averaged growth rate ⟨*g*⟩. The bottom-right corner presents the mathematical form of the sufficient condition, which depends on *β*, maximum death rate *d*_0_, minimum degree *k*_min_, maximum Laplacian eigenvalue *ω*_max_, and maximum relative growth rate *δ*_max_. Above the phase diagram, four final population densities after a finite time are shown for different spatial growth distributions selected from robust decline phase. The dashed line indicates the initial uniform population density. On the right, the corresponding spatial growth rate distributions are illustrated. **C**. Phase diagrams for two additional structures — star and complete graphs — under relatively high maximum growth rate *g*_0_ = 2(*N* −1) and low maximum death rate *d*_0_ = −1. For the complete graph, the sufficient condition becomes exact, perfectly matching the numerical boundary.

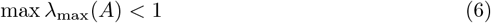

This is again a constrained optimization problem, which, as discussed earlier, reduces to selecting highly interconnected subcores (see SI for complete proof). To illustrate this framework across multi-microhabitat structures, we present migration-growth phase diagrams (*β* vs. ⟨*g*⟩) that identify parameter regimes yielding robust population decline under ⟨*g*⟩ < 0. Besides robust population decline, population responses may still vary depending on heterogeneity, resulting in a “mixed phase” region where both growth and decline are possible. Note that for any ⟨*g*⟩ < 0, the “even-spread” strategy always achieves decline, which means there’s no region where robust population growth is possible.

For simplicity, we focus on the case where *g*_0_ ≥(*N* −1) |*d*_0_|, or equivalently *c* ≥*N*. In this regime, the criterion:

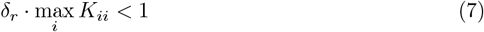

becomes an exact condition for robust population decline. Here, *δ*_*r*_ is the maximum possible relative growth rate, and max_*i*_ *K*_*ii*_ is the inverse dynamic-related centrality of the most isolated microhabitat. This condition tells us that as long as max_*i*_ *K*_*ii*_ < 1*/δ*_*r*_, we are guaranteed robust population decline. Since *K*_*ii*_ ∝ *β*^−1^ and *δ*_*r*_ ∝ ⟨*g*⟩, this condition reveals a tradeoff: higher migration suppresses growth, while higher average growth promotes persistence. To better visualize this interplay, we apply an interpolation inequality to max_*i*_ *K*_*ii*_ and rearrange terms (see SI), yielding a simplified analytical boundary for robust decline based only on structural extremes:

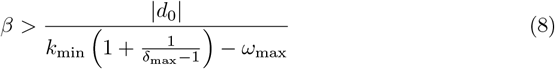

Here, *k*_min_ is the minimum degree and *ω*_max_ the largest Laplacian eigenvalue of the structure. We explicitly write *δ*_max_ = *δ*_*r*_ to emphasize its role as the extreme growth parameter. This bound clearly captures the fundamental tradeoff: to suppress populations with high intrinsic growth (*δ*_*r*_), we need stronger inter-microhabitat migration (*β*). For a complete graph, where all nonzero Laplacian eigenvalues are equal, this inequality becomes exact (see SI). In Figure 4B, we illustrate the phase diagram for a 12-node cycle graph. The blue region (top left) corresponds to robust decline (max *λ*_max_(*A*) < 1), while the white region represents the mixed phase. Our analytical bound provides a higher estimate for the true numerical boundary.

In cancer research, the star graph is especially relevant for modeling metastasis from a primary site to distant tissues [49, 51]. Similarly, in bacterial communities, fully connected networks may arise [74, 75]. Figure 4C presents phase diagrams for both star and complete graphs. In the complete graph (bottom panel), the sufficient condition precisely matches the numerical boundary.

For these two structures, we can derive closed-form expressions for max_*i*_ *K*_*ii*_. For star graph, the largest *K* is at a leaf, and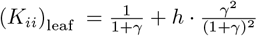, where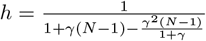. For complete graph, all nodes are identical, and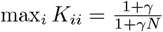. Thus, for graphs like these, the boundary line can be exactly determined by solving *δ*_*r*_ · max_*i*_ *K*_*ii*_ = 1. For more complex graphs where max_*i*_ *K*_*ii*_ is not analytically tractable (e.g., even cycles), our extreme-bound approximation from equation (8) still provides a fast and reliable method to identify regions of robust population decline.

## Discussion

Complex biological systems, such as cancer and bacterial populations, are deeply intertwined with network structure, growth dynamics, and spatial interactions. In this study, we introduced a kernel transformation that enabled the derivation of a new theoretical criterion predicting population-level responses across arbitrary multi-microhabitat structures under spatially heterogeneous drug environments. This transformation naturally links population dynamics to a centrality measure known as forest closeness centrality, which we refer to as *dynamic-related centrality* in our framework. Our findings demonstrate that increasing connectivity between microhabitats consistently promotes population decline, effectively mimicking the impact of increasing the migration rate. Furthermore, we show that the optimization of spatial drug heterogeneity for inducing population decline can be reduced to a subgraph selection problem on the transformed fully connected graph. In addition, we identify a parameter regime in the growth-migration phase diagram that supports robust population decline — a guaranteed clearance across all spatial drug configurations if the actual spatial drug heterogeneity is unknown. We derive a sufficient theoretical condition characterizing this phase, capturing the interplay between spatially averaged growth and migration rates. Notably, this condition becomes exact when the underlying microhabitat structure is a complete graph.

In [44], perturbation approximation was successfully applied to derive an analytical expression for the largest eigenvalue *λ* _max_ (*G* − *βL*), where 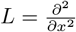 is the Laplacian operator with two absorbing boundaries on a one-dimensional continuous space. The approximated form revealed two optimized strategies for spatial drug arrangement by examining the unperturbed largest eigenvector of *L*: the “importance” of spatial positions is ranked by the squared components of this eigenvector. And it’s found that, assigning drug-free growth rates *g*_0_ to central positions tends to mitigate population decline, while assigning *g*_0_ near absorbing boundaries tends to accelerate it. However, when the spatial structure becomes discrete, and the driving force of decline arises not from absorbing boundaries, but from “sink” microhabitats with negative growth rates, perturbation theory fails to accurately capture the system’s behavior—it reflects only the effect of spatially averaged growth. In contrast, our kernel transformation introduces a new matrix *A* = *D*^1*/*2^*KD*^1*/*2^, where *K* = (*I* + *γL*)^−1^ is the regularized Laplacian kernel and *D* is the diagonal matrix of growth-related weights. Even first-order perturbation theory fails on *K*, since *K* and *L* share the same eigenbasis, and thus the largest eigenvector still assigns equal importance to all microhabitats.

Despite this, the diagonal elements *K*_*ii*_, which correspond to forest closeness centrality — or what we refer to as “dynamic-related centrality” — provide a meaningful measure for ranking the relative importance of microhabitats. Although perturbation theory is insufficient in this setting, the kernel matrix *K* still allows us to heuristically infer microhabitat importance and guide spatial drug dosing strategies. In particular, it enables a greedy subcore selection strategy on the transformed fully connected graph, as discussed in the main text. This insight allows us to extend the one-dimensional intuition: assigning *g*_0_ to central or boundary positions depends on the controllability context. In our discrete framework, the “center” corresponds to a well-interconnected core, while the “edges” represent relatively isolated subgraphs. Thus, through kernel transformation, we heuristically recover spatial clearance strategies based on dynamic-related centralities, even in cases where classical perturbation methods break down.

Across-node properties such as centrality measures play a crucial role in graph theory and network analysis. For instance, subgraph centrality quantifies node importance based on participation in subgraphs [76], with a bias toward smaller subgraphs. Interestingly, our dynamic-related centrality *C* exhibits a similar relationship with traditional closeness centrality, correlating with node degree, and other centrality measures such as Katz or eigenvalue centrality [77, 78]. Recent work on dynamic centrality has aimed to integrate both topological and dynamical properties of nodes to more accurately identify influential spreaders in static and temporal networks [79–81]. These studies emphasize the importance of incorporating temporal aspects into centrality metrics, as static measures fail to capture the evolving nature of real-world networks, crucial for predicting node importance in dynamic systems [80, 82]. Empirical studies demonstrate that these methods outperform conventional metrics across various scenarios, including information and disease spreading [81, 82]. However, previous formulations of dynamic centrality primarily focus on the addition or deletion of nodes within a network, meaning they remain fundamentally structural measures that describe the dynamics of the network itself. In contrast, our dynamic-related centrality focuses on predicting the results of dynamics occurring on the network, and it naturally emerges from the dynamics itself by doing the kernel transformation. This distinction makes our proposed measure novel, and we term it “dynamic-related centrality” to distinguish it from “dynamic centrality” for the dynamics of the network. We anticipate that this new centrality will expand existing centrality classifications and provide direct insights into dynamic properties without requiring complex simulations. Interestingly, our matrix *K*, which defines the dynamic-related centrality *C*, shares the same mathematical form as the classical regularized Laplacian kernel (RLK) [83] — a well-known graph similarity measure based on the Laplacian matrix *L*. The RLK is defined as *K* = (*I* + *γL*)^−1^, where *I* is the identity matrix, *L* is the Laplacian, and *γ >* 0 is a tunable regularization parameter that controls the global influence of the Laplacian. Despite this shared mathematical form, we emphasize important conceptual differences. First, in our growth-migration framework, the parameter 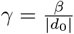 arises naturally from the underlying population dynamics, rather than being introduced as a free parameter. Second, while the RLK is typically used to analyze off-diagonal elements to quantify node-to-node similarity, our analysis centers on the diagonal entries *K*_*ii*_, which correspond to forest closeness centrality and reflect the self-centrality of each node in the dynamic context (While *λ*_max_(*A*) can be interpreted through a walk-based perspective—capturing contributions from paths traversing multiple nodes thus considering interactions between different nodes — this interpretation is beyond the scope of the present work and is not central to our theoretical framework). Establishing these connections between our dynamic-related centrality and classical graph-theoretic measures not only enhances the interpretability of our framework but also expands its potential applications in network-based modeling of biological systems.

In the main text, we did not emphasize the upper bound 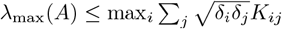, which not only serves as a useful heuristic for guiding optimized drug allocation strategies, but also reveals deeper structural insights. Specifically, we find that this maximum row sum is tightly linked to the presence of local symmetries in the system. When the matrix *A*, or a principal submatrix *A*[*S*], is circulant, the inequality becomes an equality: 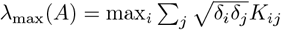. Although spatial drug heterogeneity generally breaks most global symmetries in the underlying structure, certain local symmetries may still be preserved. In such cases, this equality offers a convenient and accurate method for predicting population response by directly evaluating the maximum row sum of *A*. Detailed derivations, along with illustrative examples, are provided in the Supplementary Information.

Understanding population responses to drug treatment, such as antibiotics or chemotherapy, is critical for assessing therapeutic success. Recent studies in both bacterial and cancer systems have identified an interesting phenomenon known as “bistability”, where small differences in key parameters—such as drug concentration or growth rate—can lead to drastically different outcomes [84–88]. This concept helps explain variability in clinical outcomes. Here, we propose an alternative explanation for such bi-outcome scenarios: spatial drug heterogeneity and differences in microhabitat connectivity may also contribute to distinct clearance outcomes, independent of bistability. This insight could inform the development of new treatment strategies tailored to spatial drug distribution patterns.

Experimental studies have demonstrated the impact of spatial structure on growth [44, 89], competition, and evolution [90–93]. Our theoretical findings may aid in the interpretation of experimental results and guide the design of future studies.

Combining theoretical and experimental approaches can enhance our understanding and contribute to the development of spatially explicit drug dosing strategies.

This study explored population responses across various microhabitat structures and spatial drug heterogeneities, revealing insights beyond previous research that primarily focused on homogeneous structures and drug distributions. However, several extensions remain for future research. First, we considered only non-frequency-dependent drug-induced growth in a single cell type. Although experimental or clinical data on drug-mediated growth interactions remain scarce, some recent studies have begun to address this gap [94, 95]. Future work could incorporate frequency-dependent growth effects and multi-species interactions using evolutionary game theory [96–100] to uncover additional theoretical results and experimental comparisons. Second, we did not account for temporal drug decay or fluctuations in our models. Future work could integrate spatial-temporal drug variations based on patient-specific pharmacokinetics/pharmacodynamics (PK/PD) [25] to inform personalized treatment strategies that ensure bacterial or cancer clearance.

## Acknowledgments

This study was supported by NIH R35GM124875 (K. W.). We’d like to sincerely thank suggestions from Dr. David Lubensky and Dr. Michal Zochowoski.

## Footnotes

1 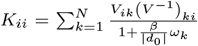, with *V* and *{ω*_*k*_*}* representing the eigenvector matrix and eigenvalues of the Laplacian matrix *L*, respectively. It contains only structure information, with maximum drug-induced death rate |*d*_0_| as the baseline growth.

