## Supplementary Information for "Deciphering and steering population-level response under spatial drug heterogeneity on microhabitat structures"

#### Supporting Information

This document contains the supporting information for the main manuscript titled "Deciphering and steering population-level response under spatial drug heterogeneity on microhabitat structures". The sections below contain additional figures, methods, and data.

#### Deterministic approximation under large population size N

Antibiotic-induced clearance of bacteria is usually stochastic[1]. For the microscopic dynamics, if we consider single cell divison rate $\lambda(D)$, lysis rate $\phi(D)$ under drug dose, and migration rate $\beta$, we have

$$X_i \xrightarrow{\lambda_i(D)} 2X_i$$

$$X_i \xrightarrow{\phi_i(D)} \varnothing_i$$

$$X_i \xrightarrow{L_{ij}\beta} X_j$$

$X_i$ represents an individual cell in ith microhabitat. Cells can only migrate from ith microhabitat to jth microhabitat if $L_{ij} = 1$. Thus it can be described by a master equation[2, 3]:

$$\frac{dP(n_i)}{d\tau} = T^+[n_i - 1]\,P(n_i-1)+T^-[n_i+1]\,P(n_i+1)-(T^+[n_i]+T^-[n_i])P(n_i).$$

$i = 1, \cdots N$. $N$ denotes the total number of microhabitats, with $T^+[n_i] = \lambda n_i - \beta\sum_j^N L_{ij}n_j$, $T^-[n_i] = \phi n_i - \beta\sum_j^N L_{ij}n_i$. If we assume that the population size is $S$, and introduce the rescaling $x_i = \frac{n_i}{S}, t = \frac{\tau}{S}, \rho(x_i, t) = SP(n_i)$, for $S \gg 1$, we can do a system size expansion or Van Kampen expansion[4, 5, 6]

$$\frac{d}{dt}\rho(x_i,t) = -\frac{d}{dx_i}[(T^+(x_i)-T^-(x_i))\rho(x_i,t)]+\frac{1}{2}\frac{d^2}{dx_i^2}\left[\frac{T^+(x_i)+T^-(x_i)}{S}\rho(x_i,t)\right],$$

with higher order terms in $S^{-1}$ neglected. Rewrite it into a Langevin equation

$$\frac{dx_i}{dt} = T^+(x_i) - T^-(x_i) + \sqrt{\frac{T^+(x_i) + T^-(x_i)}{S}}\eta$$

Let $S \to \infty$ so the second term describing stochastic effect diminishes. Replace $x_i$ with $n_i$ and we have the deterministic form

$$\frac{dn_i}{dt} = (\lambda_i - \phi_i)n_i - \beta \sum_j^N L_{ij}(n_j - n_i)$$

Rewrite $u_i = n_i$, define the effective growth rate for population $g_i = \lambda_i - \phi_i$, and write the equations in a matrix form

$$\frac{du}{dt} = -\beta L u + G u = \Omega u \tag{S1}$$

where $L$ denotes the Laplacian matrix and $G$ is a diagonal matrix with elements corresponding to bacterial growth rates, as shown in the main text. In our model system, we seek to clarify the interpretation of *large population size*. In experimental measurements of bacterial drug dose responses, bacterial populations typically exhibit exponential growth until the optical density (OD) reaches approximately 0.1 to 0.2, which corresponds to a cell density of $10^7$ to $10^8$ cells. In practical applications, when the bacterial population size $S$ exceeds $10^4$, stochastic effects can often be neglected, allowing for deterministic approximations. Notably, in clinical settings, bacterial populations generally exceed $10^4$ cells per mL by the time they become detectable in screenings or when symptoms manifest [7]. For respiratory pathogens, bacterial loads in the lungs commonly exceed $10^5$ CFU/mL [8], while a threshold of $10^5$ CFU/mL is conventionally used to diagnose urinary tract infections (UTIs) in standard urine cultures [9]. Beyond pathogenic contexts, bacterial communities on the skin also follow well-defined population densities. For example, a typical skin pore harbors approximately $5 \times 10^4$ *Cutibacterium acnes* cells [10, 11]. Given these population scales, our model considers bacterial densities in the range of $[10^4, 10^8]$, thereby justifying the deterministic approximation by neglecting stochastic fluctuations associated with bacterial growth and antibiotic clearance.

### Population decline criterion derivation by kernel transformation

Instead of directly calculating the largest eigenvalue $\lambda_0 = \|\Omega\| = \|G - \beta L\|$, for decline or clearance of the bacterial population, we focus on

$$\Omega = G - \beta L \prec 0 \tag{S2}$$

as population declines to a stable state governed by $\Omega$. We want to know, for different spatial drug heterogeneities, how this population response is tuned.

However, we cannot tune growth rates at each microhabitat arbitrarily since drug-induced growth rates have dose response limits. The diagonal entries of the growth matrix $G$, $\{g_i\}$, are bounded by $[d_0, g_0]$, the maximum death rate induced at a very high drug concentration, and the maximum growth rate when population is living in a drug-free environment, where $d_0 < 0$ and $g_0 > 0$ for bactericidal drugs like Ampcillin[12]. Do a decomposition as follows, where $\alpha > 0$ can be any absolute value between maximum death rate $d_0$ and maximum growth rate $g_0$, for example $\alpha = |d_0|$ or $\alpha = |\langle g \rangle|$. Here in this work we choose $\alpha = |d_0|$ for interpretation convenience, and this will be illustrated later. Thus we have

$$G = \alpha \cdot \begin{pmatrix} -1+\delta_1 & & & \\ & -1+\delta_2 & & \\ & & \ddots & \\ & & & -1+\delta_N \end{pmatrix} = -\alpha I + \sqrt{\alpha} D^{\frac{1}{2}} \cdot \sqrt{\alpha} D^{\frac{1}{2}}$$

where $I$ is the identity matrix, $D^{\frac{1}{2}} = \text{diag}(\sqrt{\delta_1}, \sqrt{\delta_2}, \ldots, \sqrt{\delta_N})$ is the diagonal matrix preserving growth information, and $\delta_i = 1 + \frac{g_i}{\alpha}$. Notably, $G - \beta L$ will be stable if and only if $\beta L - G$ is positive definite. We can express $\beta L - G$ in the form

$$\beta L - G = \beta L + \alpha I - \sqrt{\alpha} D^{\frac{1}{2}} \cdot I \cdot \sqrt{\alpha} D^{\frac{1}{2}}$$

Applying the Schur complement, we can see that $\beta L + \alpha I - \sqrt{\alpha} D^{\frac{1}{2}} \cdot I \cdot \sqrt{\alpha} D^{\frac{1}{2}}$ will be positive definite if and only if the matrix

$$X = \begin{pmatrix} \beta L + \alpha I & \sqrt{\alpha} D^{\frac{1}{2}} \\ \sqrt{\alpha} D^{\frac{1}{2}} & I \end{pmatrix}$$

is positive definite. Since all eigenvalues of $\beta L + \alpha I$ are strictly positive, $\beta L + \alpha I$ is invertible. By considering the other Schur complement of this block matrix, we can see that $X$ will be positive definite if and only if

$$I - \sqrt{\alpha} D^{\frac{1}{2}} (\beta L + \alpha I)^{-1} \sqrt{\alpha} D^{\frac{1}{2}} > 0 \Longleftrightarrow D^{\frac{1}{2}} (I + \frac{\beta}{\alpha} L)^{-1} D^{\frac{1}{2}} \prec I$$

Define $A = D^{\frac{1}{2}} K D^{\frac{1}{2}}$, where $K = (I + \frac{\beta}{\alpha} L)^{-1}$. Putting it all together, we see that $G - \beta L$ will be stable if and only if all the eigenvalues of $A$ is smaller than corresponding eigenvalues of $I$. Since all the eigenvalues of $I$ is equal to 1, then we require

$$\lambda_{\max}(A) < 1 \tag{S3}$$

Recall $\alpha = |d_0|$, so $\delta_i = \frac{g_i - d_0}{|d_0|} = 1 + \frac{g_i}{|d_0|}$ becomes the relative spatial derivation of growth rate compared to the maximum death rate $d_0$. This $d_0$ is only related to drug type, bacteria type and nutrition supply. Every $\delta_i$ is in the range of $[0, 1 + \frac{g_0}{|d_0|}]$. We require $\langle \delta \rangle < 1$, or equivalently $\langle g \rangle < 0$ to mimic the spatially averaged killing effect of bactericidal drug. We want to show that even the spatially averaged growth rate is negative (population decline under homogeneous growth), the population responses can be diverse, governed by the structure of $A$. By observation, we find out $K = (I + \frac{\beta}{|d_0|}L)^{-1}$ has a form of regularized Laplacian kernel[13]. This kernel function can also be seen as a discrete version of Green's functions on graphs. It's a measurement of how much mass from one diffuses to the other. $K_{ij}$ reflects the total influence that node $i$ exerts on node $j$, or the similarity between each other. The influence decays with graph distance and degree. We can also see that $K$ is dense even if $L$ is sparse — it captures global structure from local transitions. $A = D^{\frac{1}{2}}KD^{\frac{1}{2}}$ can be interpreted as a new diffusion kernel function weighted by relative growth values. Besides $K_{ij}$, the diagonal entries $K_{ii}$ has often been used to define new centralities. $K = (I + \frac{\beta}{|d_0|}L)^{-1}$ has also been named as forest matrix[14], to define a new centrality called forest closeness centrality, in the mathematical form $C_i = \frac{N}{K_{ii}+tr(K)-2}$[15, 16]. Closeness centrality describes how close one microhabitat is to the others, indicating its ability to quickly reach others. Compared to traditional closeness centrality, this forest closeness centrality considers not only the shortest path between microhabitat $i$ and $j$, but all paths connecting them. It has the power to describe the global effect (effectively transforming the original structure to a fully-connected graph), even for disconnected graphs and isolated microhabitats.

For $A_{ij}$, we have

$$\begin{aligned}
A_{ij} &= \left(D^{\frac{1}{2}}KD^{\frac{1}{2}}\right)_{ij} \\
&= D_{ii}^{\frac{1}{2}}K_{ij}D_{jj}^{\frac{1}{2}} \\
&= \sqrt{\delta_i\delta_j}K_{ij} \\
&= \sqrt{\delta_i\delta_j}(I + \frac{\beta}{|d_0|}L)_{ij}^{-1} \\
&= \sqrt{\delta_i\delta_j}\sum_{k=1}^{N}\frac{V_{ik}\left(V^{-1}\right)_{kj}}{1+\frac{\beta}{|d_0|}\omega_k}
\end{aligned}$$

The second equality holds because $D^{\frac{1}{2}}$ is a diagonal matrix[1]. The last equality holds due to the similarity transformation of $K$[2], where $w_k$ is the kth eigenvalue of $L$, $V$ is the eigenvector matrix of Laplacian matrix $L$. $A_{ij}$ is generally nonzero and locally captures the different effects a node $i$ has on a node $j$, based on diffusion and local growth. So here $A$ describes an effective fully connected graph(clique) with different weighted edges $A_{ij}$.

**Special condition: undirected Laplacian matrix** Specifically for undirected Laplacian matrix $L$ (always with complete migration feedback), the eigen-

[1]See the SI section "Supplementary derivations" for complete proof.

[2]$K = (I + \frac{\beta}{\alpha}L)^{-1} = V\mathcal{W}V^{-1}$. $\mathcal{W}$ is the diagonal eigenvalue matrix of Laplacian matrix $L$.

vector is orthogonal because $L$ is symmetric. So $V^{-1} = V^T$. So we have

$$A_{ij} = \sqrt{\delta_i \delta_j} \sum_{k=1}^{N} \frac{v_{k,(i)} v_{k,(j)}}{1 + \frac{\beta}{|d_0|} \omega_k}, i = 1, 2, \ldots, N \tag{S4}$$

where $\sum_{k=1}^{N} v_{k,(i)}^2 = 1$ since $V$ is orthonormal.

### Derivations for bounds of $\lambda_{\max}(A)$ predicting population response

Although it's hard to derive the exact analytical form of $\lambda_{\max}(A)$ due to its inhomogeneous nature, we can instead find it's lower and upper bounds in relatively simple forms

$$\max_i K_{ii}\, \delta_i \leq \lambda_{\max}(A) \leq \max_i \sum_j \sqrt{\delta_i \delta_j}\, K_{ij} \tag{S5}$$

**Upper bound derivation** Here we introduce a new expression for the largest eigenvalue of a matrix. The dominant eigenvalue of $A$ can be rewritten as a limiting norm

$$\max \lambda(A) = \lim_{k \to \infty} \left\| A^k \right\|^{1/k}$$

where $\| \cdot \|$ is any norm[17]. For any finite positive integer $k$, we always have

$$\max \lambda(A) = \lim_{k \to \infty} \left\| A^k \right\|^{1/k} \leq \left\| A^k \right\|^{1/k}.$$

This induces an upper bound for the largest eigenvalue of matrix $A$. Here we choose $\| \cdot \|_\infty$, the largest row sum of absolute value of a matrix, as our norm $\| \cdot \|$, and let $k = 1$, we get

$$\begin{aligned} \max \lambda(A) &\leq \|A\|_\infty \\ &= \max \left( |A_{ii}| + \sum_{j, i \neq j} |A_{ij}| \right) \\ &= \max \left( |\delta_i K_{ii}| + \sum_{j, i \neq j} |\sqrt{\delta_i \delta_j} K_{ij}| \right) \\ &= \max_i \sum_j \sqrt{\delta_i \delta_j} K_{ij} \end{aligned}$$

The last equality holds since $\delta_i \geq 0, K_{ii} > 0, K_{ij} > 0$[3]. Thus we derive the upper bound for $\lambda_{\max}(A)$. If $\max_i \sum_j \sqrt{\delta_i \delta_j} K_{ij} < 1$, we have $\lambda_{\max}(A) \leq$

[3] For $K_{ii} > 0, K_{ij} > 0$, see the SI section "Supplementary derivations" for complete proof.

$\max_i \sum_j \sqrt{\delta_i\delta_j}K_{ij} < 1$, inducing population decline. Thus $\max_i \sum_j \sqrt{\delta_i\delta_j}K_{ij} < 1$ can serve as a sufficient criterion to predict population decline, or a necessary criterion for population growth.

This first-order sufficient condition for population decline can also be obtained by applying Gershgorin's Circle Theorem for the dominant eigenvalue[4]. Since $K_{ii} > K_{ij}$[5], the further the distance between microhabitat i and j, the more the interaction effect $\sqrt{\delta_i\delta_j}$ is decayed by $K_{ij}$. We can tighten this bound $\|A^k\|^{1/k}$ by considering $k \geq 2$, which approaches the largest eigenvalue of $A$ from above as $k \to \infty$.

**Walk-based interpretation of $\lambda_{\max}(A) = \lim_{k\to\infty} \|A^k\|^{1/k}$ and cluster expansion** Let $\mathcal{W}_k(i \to j)$ be the set of all length-$k$ walks from node $i$ to $j$, then:

$$(A^k)_{ij} = \sum_{\omega \in \mathcal{W}_k(i\to j)} \text{weight}(\omega)$$

$$\lambda_{\max}(A) = \lim_{k\to\infty} \left( \max_i \sum_j \sum_{\omega \in \mathcal{W}_k(i\to j)} \text{weight}(\omega) \right)^{1/k}$$

For example, we can have $\text{weight}(\omega) = A_{il_1}A_{l_1l_2}\cdots A_{l_{k-2}l_{k-1}}A_{l_{k-1}j}$. This is like a cluster expansion, and in sparse graphs with localized growth, the expansion truncates early much like in quantum field theory where only low-order diagrams contribute.

**Lower centrality-based bound derivation** Here we prove the lower bound $\max_i K_{ii}\,\delta_i \leq \lambda_{\max}(A)$ by leveraging the walk-based interpretation. If we denote the row with largest row sum for $A^k$ as $i^*$, then we have

$$\begin{aligned}
\lambda_{\max}(A) &= \lim_{k\to\infty} \left( \max_{i^*} \sum_j \sum_{\omega \in \mathcal{W}_k(i^*\to j)} \text{weight}(\omega) \right)^{1/k} \\
&= \lim_{k\to\infty} \left( \max_{i^*} \left( A^k_{i^*i^*} + \sum_{\omega \in \mathcal{W}_k(i^*\to i^*)\setminus(i^*)^k} \text{weight}(\omega) + \sum_{j, j\neq i^*} \sum_{\omega \in \mathcal{W}_k(i^*\to j)} \text{weight}(\omega) \right) \right)^{1/k} \\
&\geq \lim_{k\to\infty} \left( \max_i A^k_{ii} + \left( \sum_{\omega \in \mathcal{W}_k(i\to i)\setminus(i)^k} \text{weight}(\omega) + \sum_{j, j\neq i} \sum_{\omega \in \mathcal{W}_k(i\to j)} \text{weight}(\omega) \right) \right)^{1/k} \\
&\geq \lim_{k\to\infty} \left( \max_i A^k_{ii} \right)^{1/k} \\
&= \max_i A_{ii} \\
&= \max_i K_{ii}\delta_i.
\end{aligned}$$

[4]Also see the SI section "Supplementary derivations" for complete proof.

[5]For $K_{ii} > K_{ij}$, see the SI section "Supplementary derivations" for complete proof.

The second equality decomposes the $k$-step self-walk at node $i$ into three parts: repeated self-walks at $i$, walks that start and end at $i^*$ in $k$ steps, and all other walks originating from $i^*$ that reach different microhabitats. The third inequality holds since we can choose a new row $i$ which contains $\max_i A_{ii}^k$ instead of the maximum row sum. It can be equal when $i = i^*$ - the maximum row sum contains the maximum diagonal entry. The fourth inequality holds because all weights are non-negative. Thus we have proved $\max_i\ K_{ii}\,\delta_i \leq \lambda_{\max}(A)$.

Similarly, if $\max_i K_{ii}\delta_i > 1$, we have $\lambda_{\max}(A) \geq \max_i K_{ii}\delta_i > 1$, inducing population growth. Thus $\max_i K_{ii}\delta_i > 1$ can serve as a sufficient criterion to predict population growth, or a necessary criterion for population decline. Since $K_{ii} \propto C_i^{-1}$ as the inversed dynamic-related centrality, here we call this lower bound the centrality-based bound. Since this lower bound has a simpler form than the upper bound, in practice, we can always perform a quick diagnostic check of the population response using the quantity $\max_i K_{ii}\,\delta_i$ first. If $\max_i K_{ii}\,\delta_i > 1$, we can immediately predict population growth. Conversely, if $\max_i K_{ii}\,\delta_i < 1$, this provides a necessary condition for population decline. As a second step, we evaluate the upper bound $\max_i \sum_j \sqrt{\delta_i \delta_j}\, K_{ij}$; if this quantity is also smaller than 1, we can conclude that population decline will occur.

This zero-order lower bound can also be obtained by applying the Rayleigh quotient for the dominant eigenvalue[6]. We can tighten this lower bound by using power iteration for the Ryleigh quotient, or considering more weights of walks in $\lim_{k\to\infty} \left\|A^k\right\|^{1/k}$.

**Why not directly apply bound derivations to the largest eigenvalue of original matrix $\lambda_{\max}(G-\beta L)$?** If we apply the same lower and upper bounds to $\lambda_{\max}(G-\beta L)$, we have $\max_i(G-\beta L)_{ii} = \max_i(g_i - \beta k(i)), \max_i \sum_j (G-\beta L)_{ij} = \max_i(g_i - k(i) + k(i)) = \max_i g_i$, where $k(i)$ is the degree of the ith microhabitat. Thus $\max_i(g_i - \beta k(i)) \leq \lambda_{\max}(G-\beta L) \leq \max_i g_i$. The lower bound only captures local degree information, and the upper bound gives no information since it's already well-known that $\lambda_{\max}(G-\beta L)$ can be no larger than the largest growth rate.

### Special cases when lower or upper bounds become exact

Although $A$ describes an effective fully connected graph, for some $\delta_i = 0$, the edges can be effectively removed, and the substructures will become smaller cliques, represented by $A_S$, where $S$ denotes the remaining number of vertices. For the lower centrality-based bound, $\lambda_{\max}(A_S) = \max_i K_{ii}\delta_i$ when only 1 microhabitat has non-zero relative growth $\delta_i$. And $\max_i K_{ii}\delta_i < 1$ becomes an exact condition for predicting population decline. For the upper bound, $\lambda_{\max}(A_S) = \max_i \sum \sqrt{\delta_i\delta_j} K[S]_{ij}$ when $A_S$ is a circulant matrix preserving the local translational symmetry of the original graph structure. And

[6]Also see the SI section "Supplementary derivations" for complete proof.

$\max_i \sum \sqrt{\delta_i \delta_j} K[S]_{ij} < 1$ becomes an exact condition for predicting population decline too. Here we will discuss the upper bound first, since $\lambda_{\max}(A_S) = \max_i K_{ii}\delta_i$ can also be seen as a special case of $\lambda_{\max}(A_S) = \max_i \sum \sqrt{\delta_i \delta_j} K[S]_{ij}$.

#### $\lambda_{\max}(A_S) = \max_i \sum \sqrt{\delta_i \delta_j} K[S]_{ij}$: preserved local symmetry

Here we'd like to prove $\lambda_{\max}(A_S) = \max_i \sum \sqrt{\delta_i \delta_j} K[S]_{ij}$ under local symmetry by the following formalized theorem.

**Theorem 1** (Exact via local symmetry)**.** *Let $A = D^{1/2} K D^{1/2} \in \mathbb{R}^{n\times n}$, where:*

- *$K = (I + \gamma L)^{-1}$ is the regularized Laplacian kernel of a graph $G$,*
- *$D = \mathrm{diag}(\delta_1, \ldots, \delta_n)$ is a diagonal matrix with $\delta_i \geq 0$,*

*and let $S \subseteq V$ denote the set of active nodes (i.e., $\delta_i > 0$). Define the restricted matrix*

$$A_S := D_S^{1/2} K[S] D_S^{1/2},$$

*where $D_S = \mathrm{diag}(\delta_i)_{i\in S}$ and $K[S]$ is the submatrix of $K$ indexed by $S$.*

*Suppose $A_S$ is a circulant matrix, or the following conditions hold:*

1. *The induced subgraph on $S$ is symmetric under a nontrivial subgroup of automorphisms of $G$,*
2. *The growth vector $\delta_S$ is invariant under this symmetry,*
3. *The matrix $A_S$ has identical rows, i.e., $A_S \mathbf{1} = q\mathbf{1}$.*

*Then for all integers $k \geq 1$, the walk-based estimator of largest eigenvalue*

$$\lambda_k := \left( \max_{i\in S} \sum_{j\in S} (A_S^k)_{ij} \right)^{1/k}$$

*is exact, i.e.,*

$$\lambda_k = \lambda_{\max}(A_S) = \lambda_{\max}(A).$$

*In particular, the largest eigenvalue can be achieved at $k = 1$:*

$$\lambda_1 = \max_{i\in S} \sum_{j\in S} (A_S)_{ij} = \max_{i\in S} \sum_{j\in S} \sqrt{\delta_i \delta_j} K[S]_{ij} = \lambda_{\max}(A).$$

**Walk-based Proof** According to the walk-based interpretation, we can rewrite the dominant eigenvalue of $A$ as $\lambda_{\max}(A) = \lim_{k\to\infty} \left( \max_i \sum_j \left(A^k\right)_{ij} \right)^{1/k}$. For those nodes with $\delta_i = 0$, thus we have $A_{ij} = 0$, where $i = S+1, \cdots, N$, and $j = 1, \cdots N$. Since $(A^k)_{ij}$ represents the total weight of all walks of length $k$ from node $i$ to node $j$, setting $A_{ij} = 0$ effectively eliminates any walk that includes the edge $(i, j)$, thereby restricting the remaining walk contributions to those entirely within the S-clique. Thus we have $\lambda_{\max}(A) = \lim_{k\to\infty}\left(\max_i \sum_j \left(A^k\right)_{ij}\right)^{1/k} = \lim_{k\to\infty}\left(\max_i \sum_j \left(A_S^k\right)_{ij}\right)^{1/k} = \lambda_{\max}(A_S)$.

The theorem above can be simplified as: for a circulant matrix $A_S = \operatorname{circ}(a_1, \ldots, a_S)$, we require

$$\lambda_{\max}(A_S) = \lim_{k\to\infty}\left(\max_i \sum_j \left(A_S^k\right)_{ij}\right)^{1/k} = \sum_{i=1}^{S} a_i = q$$

From walk-based interpretation, we know that the total weight of all remaining walks of length $k$ starting at node $i$ is

$$\sum_j \left(A_S^k\right)_{ij} = \sum_{\text{walks of length } k \text{ from } i} w(\omega)$$

But since $A_S$ is circulant, all nodes are symmetric. So the total weight of all walks of length $k$ from any node is the same. Hence:

$$\sum_j \left(A^k\right)_{ij} = \sum_{\omega\in\Omega_k} w(\omega) \quad \text{independent of } i$$

Each walk corresponds to an ordered product of $k$ steps from the multiset $\{a_1, \ldots, a_S\}$ The total weight of all such walks is

$$\sum_{\omega\in\Omega_k} w(\omega) = \left(\sum_{i=1}^{S} a_i\right)^k$$

The equality holds by multinomial expansion. Therefore

$$\sum_j \left(A_S^k\right)_{ij} = \left(\sum_{i=1}^{S} a_i\right)^k = q^k$$

Then:

$$\lambda_k := \left(\max_i \sum_j \left(A_S^k\right)_{ij}\right)^{1/k} = \left(q^k\right)^{1/k} = r \Rightarrow \lim_{k\to\infty} \lambda_k = q = \sum_{i=1}^{S} a_i$$

This equals $\lambda_{\max}(A)$ as expected.

This result provides a neat analytical expression for $\lambda_{\max}(A)$ under preserved local symmetry. It also reveals that when symmetry is broken — either by structural heterogeneity or uneven growth distributions — the quantity $\max_i \sum_j \sqrt{\delta_i\delta_j} K[S]_{ij}$, which captures the contribution from one-step walks (i.e., direct interactions), deviates from $\lambda_{\max}(A)$ and instead serves as an upper bound.

**special case: active subset of $\delta_i = \delta$** For some nodes with local symmetry in the original graph, let their relative growth $\delta_i = \delta$. Then the exact condition of population decline becomes $\delta \sum_{j \in S} K[S]_{ij} = \delta q_S < 1$, where $q_S$ is the constant value of the row sum of the sub-matrix $K[S]$.

Let's give an interesting example. Consider an 8-node cycle graph $C_8$ with 2 different relative growth distributions. For case I, we have $\{\delta_i\} = \{0, 0, 0, 0, 0, 0, \delta, \delta\}$, with active nodes 6 and 7 (adjacent). For case II, we have $\{\delta_i\} = \{0, 0, 0, \delta, 0, 0, 0, \delta\}$, with active nodes 3 and 7 (opposite). So $\lambda_I(A) = \delta(K_{11} + K_{12})$, and $\lambda_{II}(A) = \delta(K_{11} + K_{14})$. If we let $\delta = 2, \beta = 1, g_0 = 1, d_0 = -1$, we can find out $\lambda_I(A) > 1$ and $\lambda_I(A) < 1$. So one gives us population growth, but the other induces population decline. The key reason lies in the inequality $K_{14} < K_{12}$, which reflects the decay with distance inherent in the regularized Laplacian kernel. This explains why a large, spatially aggregated region with positive growth tends to induce a growth response, whereas small, scattered regions with positive growth are less effective. Similar spatial response patterns have been observed in population dynamics under drug heterogeneity in a continuous one-dimensional space [18].

When the set $\{\delta_i\}$ is not evenly distributed, a small remaining submatrix $A_S$ (e.g., for $S = 2$ or $S = 3$) corresponds to a small motif, such as a two-microhabitat structure or a simple triangle. In such cases, the effective network is still analytically tractable, and the largest eigenvalue $\lambda_{\max}(A_S)$ can be computed explicitly using known formulas, although it no longer equals $\max_i \sum_{j \in S} \sqrt{\delta_i \delta_j} K_{ij}$.

#### $\max_i K_{ii}\delta_i < 1$ becomes an exact condition when $\mathrm{rank}(A) = 1$

We observe that since $A = D^{\frac{1}{2}} K D^{\frac{1}{2}}$, and $K$ is invertible, the rank of $A$ is determined by the rank of $D^{\frac{1}{2}}$, i.e.,

$$\mathrm{rank}(A) = \mathrm{rank}(D^{\frac{1}{2}}).$$

If $\mathrm{rank}(A) = 1$, it follows that $\mathrm{rank}(D^{\frac{1}{2}}) = 1$, which corresponds to a special case where the growth rate distribution is given by $\{\delta_i\} = \{\delta_1, 0, 0, \ldots, 0\}$.

To examine this scenario, consider a growth rate distribution in which only a single microhabitat exhibits a positive growth rate, while all others are assigned the maximum death rate. This setup eliminates interaction effects between different microhabitats, allowing us to isolate the contribution from the lone microhabitat with positive growth. Under this condition, the upper bound $\lambda_{\max}(A) \le \max_i \sum_j \sqrt{\delta_i \delta_j} K_{ij}$ simplifies to

$$\lambda_{\max}(A) \le \max_i \delta_i K_{ii},$$

since all other $\delta_j$ values are zero. Combined with the previously established lower bound,

$$\lambda_{\max}(A) \ge \max_i \delta_i K_{ii},$$

we obtain an exact expression:

$$\lambda_{\max}(A) = \max_i \delta_i K_{ii} = \delta_1 K_{11}.$$

##### When matrix $K$ is approximately diagonal

Another way to make either the lower or upper bound become exact is to ensure that the interaction terms $\sum_{j\neq i} \sqrt{\delta_i \delta_j} K_{ij}$ are negligible. This can be achieved by enforcing either $\delta_j \to 0$ or $K_{ij} \to 0$. Accordingly, we identify two conditions:

**a. Growth rate distribution:** If enough growth rates $g_i$ are close to the baseline death rate $d_0$, then $\delta_i \to 0$ for many nodes. In the extreme case where all $\delta_i = 0$, the condition becomes exact with $\lambda_{\max}(A) = 0$.

**b. Migration or connectivity:** If either the migration rate $\beta$ or the connectivity (e.g., degree $k(i)$) becomes sufficiently small, the nodes become nearly isolated. In the limit of complete isolation, all off-diagonal $K_{ij} \to 0$, and the sufficient condition becomes exact.

#### Approximated inverse relationship between $K_{ii}$, migration rate $\beta$, averaged edge number $|E|$

Understanding the relationship between dynamic-related centrality $C$ and dynamical or structural properties of the network, such as the migration rate $\beta$ and the number of edges $|E|$, is crucial for characterizing population decline. In particular, we employ an approximation method to roughly analyze how centrality $C$ scales with increasing migration rate and increasing edge density.

##### Empirical $\max_i K_{ii}\delta_i - \beta$ relationship and analytical appoximation

**Empirical result: $\max K_{ii}\delta_i$ decreases as $\beta$ increases** Migration plays a crucial role in shaping population dynamics by redistributing individuals across spatially structured environments. As shown in the main text and Figure S1, the $K_{ii}$ exhibits an inverse relationship with migration rate $\beta$ under the same spatial drug distribution, similar to its dependence on edge number $|E|$. This relationship emerges from the mathematical formulation of $K_{ii}$, where increased migration weakens the influence of local growth variations and leads to a more homogenized system.

The transition between population growth and decline can be captured through the behavior of $\max K_{ii}\delta_i$, which serves as a criterion for assessing system-wide viability. As migration rate $\beta$ increases, the maximum $\max K_{ii}\delta_i$ value decreases, indicating a diminishing capacity for localized population persistence. When migration is low ($\beta \approx 0$), populations can sustain themselves through spatial heterogeneity. However, as migration increases, the diffusive spreading of individuals reduces local advantages, eventually pushing the system into a state where decline dominates. This effect parallels the impact of increasing network connectivity, as both factors promote diffusion across the system. The results suggest that higher migration rates reduce the potential for local refuges, making populations more susceptible to global decline, a mechanism that has implications for disease spreading, bacterial resistance evolution, and ecological metapopulations.

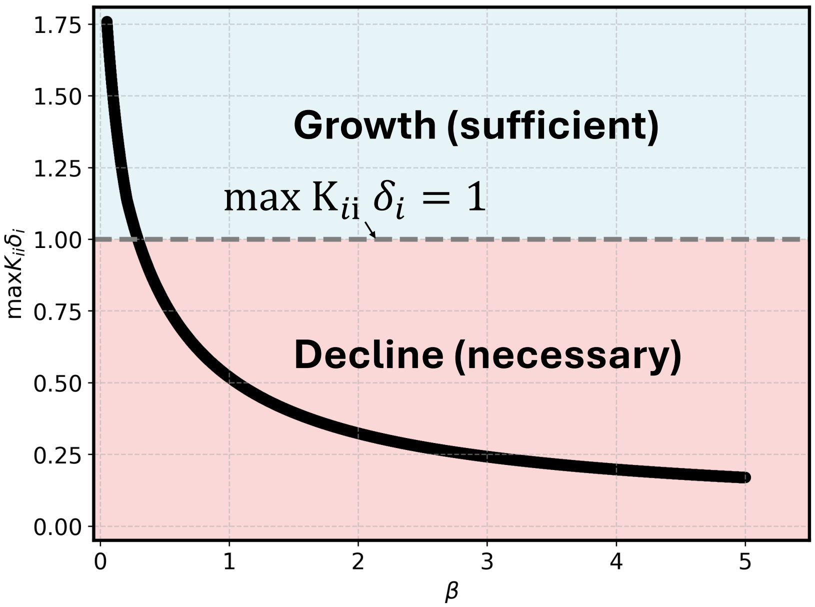


Figure S1: **Transition between growth and decline as a function of migration rate $\beta$.** The maximum $K_{ii}\delta_i$ value decreases as migration rate increases. As the curve crosses the critical threshold $\max K_{ii}\delta_i = 1$, population response shifts from growth to decline. The effect of migration is analogous to increasing network connectivity, leading to homogenization and reducing the persistence of localized populations. For the parameters, $N = 40, g_0 = 2, d_0 = -1, \langle g \rangle = -0.25$, and $\{g_i\}$ is generated by a random seed 42.

**Analyical approximation of $K_{ii} \propto \beta^{-1}$** We first consider the effect of migration rate $\beta$ on growth-migration centrality. The denominator in the inversed centrality $K_{ii}$ definition,

$$1 + \frac{\beta}{|\langle g \rangle|}\omega_k,$$

scales directly with $\beta$, meaning that as $\beta$ increases, the term $\frac{\beta}{|\langle g \rangle|}\omega_k$ grows larger, making the denominator larger for each $k$. Since the numerator contains $v_{k,(i)}^2$, which remains unaffected, the contribution of each term in the sum decreases as:

$$\frac{v_{k,(i)}^2}{1+\frac{\beta}{|\langle g\rangle|}\omega_k} \propto \frac{1}{\beta}, \quad \text{for sufficiently large } \beta.$$

Thus, for high migration rates, the total contribution to $K_{ii}$ simplifies to:

$$K_{ii} \approx \sum_{k=1}^{N} \frac{v_{k,(i)}^2}{\frac{\beta}{|\langle g\rangle|}\omega_k} = \frac{|\langle g\rangle|}{\beta} \sum_{k=1}^{N} \frac{v_{k,(i)}^2}{\omega_k}.$$

This result confirms that $K_{ii}$ approximately follows an inverse scaling law with respect to the migration rate,

$$K_{ii} \propto \beta^{-1}.$$

As a consequence, increasing $\beta$ enhances diffusion across the network, causing growth to spread more uniformly, which in turn reduces the influence of individual microhabitats.

#### Approximation of $K_{ii} \propto |E|^{-0.5}$

Consider the case of a "near-complete graph", where the network is highly connected. The inversed centrality $K_{ii}$ is given by:

$$K_{ii} = \sum_{k=1}^{N} \frac{v_{k,(i)}^2}{1+\frac{\beta}{|\langle g\rangle|}\omega_k}.$$

To understand the dependence on $|E|$, we examine the scaling behavior of the eigenvalues and eigenvectors of the Laplacian. In a nearly complete graph, the eigenvalues scale as:

$$\omega_k \propto |E|^{0.5},$$

while the eigenvector components satisfy:

$$v_{k,(i)} \propto |E|^{-0.5}.$$

Substituting these scalings into the centrality expression, the denominator for $k \geq 2$ behaves as:

$$1+\frac{\beta}{|\langle g\rangle|}\omega_k \propto |E|^{0.5},$$

and since the squared eigenvector components satisfy:

$$v_{k,(i)}^2 \propto |E|^{-1},$$

we find that the contribution to $K_{ii}$ from each term in the summation is:

$$\frac{v_{k,(i)}^2}{1 + \frac{\beta}{|\langle g \rangle|}\omega_k} \propto |E|^{-1.5}.$$

Summing over all $k$, we approximate the total contribution to $K_{ii}$ as:

$$K_{ii} \approx \frac{1}{N} + \sum_{k=2}^{N} |E|^{-1.5}.$$

Using the relation $N \approx |E|^{0.5}$, we obtain:

$$K_{ii} \approx |E|^{-0.5} + |E|^{-1}.$$

For large $|E|$, the dominant term dictates the overall scaling behavior,

$$K_{ii} \propto |E|^{-0.5}.$$

Thus, as the number of edges increases, inversed centrality $K_{ii}$ decreases, revealing an inverse relationship between $K_{ii}$ and $|E|$. This scaling suggests that in highly connected networks, centrality values become more homogeneous, as the influence of individual nodes diminishes. Beyond near-complete graphs, this inverse relationship also holds in Erdős–Rényi random graphs.

These results collectively illustrate that inversed centrality $K_{ii}$ decreases both with increasing migration rate $\beta$ and increasing edge number $|E|$. The underlying mechanism in both cases is the enhanced diffusion of growth potential across the network, which leads to a more uniform distribution and weakens the relative influence of any single node. Although the inverse relationships for both parameters are derived through approximations, and the exact scaling exponent may vary, these initial results provide an important theoretical foundation for understanding how network connectivity and migration dynamics shape population decline criteria.

### Optimized clearance strategy - a constrained optimization problem

Different spatial drug heterogeneities induce distinct population responses. To allow meaningful comparison across scenarios, we control the spatially averaged drug dose - or equivalently, the spatially averaged growth rate $\langle g \rangle$ - in our minimal model. We consider two major cases depending on whether spatial drug heterogeneity is controllable: 1. Fully controllable heterogeneity: Clinically, if we are fortunate enough to precisely control drug concentration at each microhabitat (or node), population decline can be achieved by selecting the spatial drug assignment that minimizes the largest eigenvalue - we simply find when $\min \lambda_{\max}(A) < 1$. This gives us the optimal spatial drug configuration among all possible combinations. 2. Uncontrollable or unknown heterogeneity: If spatial drug assignment cannot be precisely controlled or is unknown, we must ensure population decline under the worst-case spatial drug configuration. This requires $\max \lambda_{\max}(A) < 1$, ensuring a robust population decline that is independent of the specific drug distribution.

**Derivation of "1-remainder" strategy with uncontrollable heterogeneity, or partial controllable microhabitats** Reformalize the constrained optimization problem as follows,

$$\max_{\boldsymbol{\delta} \in [0,c]^N, \langle \delta \rangle = \frac{1}{N}\sum_i \delta_i} \lambda_{\max}\left(D^{1/2} K D^{1/2}\right)$$

where $c = 1 + \frac{g_0}{|d_0|}$ is the maximum possible relative growth value, $\boldsymbol{\delta} = (\delta_1, \ldots, \delta_N) \in [0,c]^N$ are our decision variables, $\langle \delta \rangle = \frac{1}{N}\sum_i \delta_i$ is the averaged-dose constraint. We need to prove that "1-remainder" strategy is always the optimized strategy for such a problem - $S-1$ number of microhabitats with $\delta_i = c$, $N-(S-1)$ number of microhabitats with $\delta_i = 0$, and at most 1 microhabitat with $\delta_i = \delta_r$. Here $S = \left\lceil \frac{\langle \delta \rangle}{c} \right\rceil, \quad \delta_r = N\langle \delta \rangle - (S-1)c \in [0,c]$. Although this "1-remainder" strategy cannot directly tell us the selection of subcores, it serves as a necessary condition for this constrained optimization.

Define $A(\boldsymbol{\delta}) = D^{1/2} K D^{1/2}$, since $D = \operatorname{diag}(\boldsymbol{\delta})$. Rewrite $\Lambda(\boldsymbol{\delta}) = \lambda_{\max}(A(\boldsymbol{\delta}))$ as a function of decision variable $\boldsymbol{\delta}$. Write the dominant eigenvalue in maximized Rayleigh quotient form and we get:

$$\Lambda(\boldsymbol{\delta}) = \max_{\|x\|_2 = 1} x^\top A(\boldsymbol{\delta}) x$$

Fix $\boldsymbol{\delta}$ and let $x^* = v(\boldsymbol{\delta})$ achieve the max, where $v = v(\boldsymbol{\delta}) \succ 0$ is the Perron vector, and $\|v\|_2 = 1$. By the envelope theorem,

$$\frac{\partial \Lambda}{\partial \delta_i} = \frac{\partial}{\partial \delta_i}\left(x^\top A(\boldsymbol{\delta}) x\right)\Big|_{x=x^*} = v^\top \left(\frac{\partial A}{\partial \delta_i}\right) v$$

and $\frac{\partial A}{\partial \delta_i} = \frac{\partial D^{1/2} K D^{1/2}}{\partial \delta_i} = \frac{1}{2\sqrt{\delta_i}}\left(e_i e_i^\top K D^{1/2} + D^{1/2} K e_i e_i^\top\right)$. Because $K$ is symmetric and $v$ is real, we can calcuate the first derivative $\frac{\partial \Lambda}{\partial \delta_i} = \frac{v_i^2}{2\sqrt{\delta_i}}[Kv]_i$. Define $\phi_i(\delta_i) := \frac{\partial \Lambda}{\partial \delta_i}$ for symbol convenience. For fixed $v$ and $Kv$ (they change only infinitesimally when we vary a single $\delta_i$), we find out

$$\phi_i(\delta_i) = \frac{C_i}{\sqrt{\delta_i}}, \quad C_i := \frac{1}{2} v_i^2 [Kv]_i > 0$$

Hence for $0 < \delta_i < c$, $\phi_i(\delta_i)$ is always positive, and is strictly decreasing in each coordinate. Let's introduce KKT conditions, with a multiplier $\mu$ for the average-dose constraint and box multipliers $\eta_i^-, \eta_i^+ \geq 0$ for $\delta_i \geq 0, \delta_i \leq c$. And stationarity gives $\phi_i(\delta_i) - \mu + \eta_i^- - \eta_i^+ = 0$, while complementary slackness gives us $\eta_i^- \delta_i = 0, \quad \eta_i^+ (c - \delta_i) = 0$. These conditions can tell us the status of $\phi_i(\delta_i)$ with different $\delta_i$. Below is a summary table,

| Status of $\boldsymbol{\delta_i}$ | Multiplier Conditions and Result |
|---|---|
| $0 < \delta_i < c$ | $\eta_i^- = \eta_i^+ = 0 \Rightarrow \phi_i(\delta_i) = \mu$ |
| $\delta_i = 0$ | $\eta_i^- \geq 0,\ \eta_i^+ = 0 \Rightarrow \phi_i(\delta_i) \leq \mu$ |
| $\delta_i = c$ | $\eta_i^- = 0,\ \eta_i^+ \geq 0 \Rightarrow \phi_i(\delta_i) \geq \mu$ |

Table 1: KKT conditions and resulting constraints on $\phi_i(\delta_i)$ based on $\delta_i$ bounds.

From Table 1, we see that to maximize the dominant eigenvalue, we aim to assign as many $\delta_i = c$ as possible. This yields a maximum of $\left\lfloor \frac{\langle\delta\rangle}{c} \right\rfloor$ such microhabitats, which we denote as $S-1$. There may be at most one remainder term, $\delta_r = n\langle\delta\rangle - (S-1)c \in [0, c]$, and the remaining $N-S$ microhabitats are assigned $\delta_i = 0$.

Once the $N-S$ microhabitats with zero relative growth are determined, we can restrict our analysis to the subgraph composed of nodes with non-zero $\delta_i$. This demonstrates that optimizing spatial drug heterogeneity for inducing population decline reduces to a subgraph selection problem on the transformed fully connected graph.

**Derivation of "even-spread" strategy with fully controllable heterogeneity** Here we can reformalize the constrained optimization problem as follows,

$$\min_{\boldsymbol{\delta}\in[0,c]^N,\langle\delta\rangle=\frac{1}{N}\sum_i\delta_i} \lambda_{\max}\left(D^{1/2}KD^{1/2}\right)$$

Since our average-dose constraint $\langle\delta\rangle = \frac{1}{N}\sum_i \delta_i$ is convex, and the dominant eigenvalue is also convex, this minimization problem is a classic convex optimization problem - the optimized solution will be an interior point with every possible $0 < \delta_i < c$. And now we require $\phi_i(\delta_i) = \mu$, for every microhabitat. This is usually hard to derive $\{\delta_i\}$ analytically. Since $\langle g\rangle < 0$, "even-spread" strategy with every $\delta_i = \langle\delta\rangle < 1$ is a good enough strategy to induce population decline. If under some conditions we have matrix $K$ as a strictly diagonal dominant matrix, we can have $\phi_i(\delta_i) \approx K_{ii}\delta_i = \mu$, so the optimal strategy will be approximately $\delta_i \propto (K_{ii})^{-1}$. Roughly $K_{ii}^{-1} \propto k(i)$, the degree of microhabitat. And thus the optimal strategy becomes $\delta_i \propto k(i)$, where the optimal relative growth rate of each node is made proportional to its degree. This is exactly the optimal strategy to assign curing rate and suppress epidemic outbreak in susceptible-infectious-susceptible (SIS) network dynamics[19, 20, 21, 22].

### Phase boundary of robust population decline

As mentioned in maintext, in our study, for simplicity, we consider when $g_0 \geq (N-1)|d_0|$ $(c \geq N)$. And under this case $\lambda_{\max}(A) = \max_i K_{ii}\delta_i$. To numerically determine the phase boundary for robust population decline, we consider the following constrained optimization problem

$$\max_{\boldsymbol{\delta}\in[0,c]^N, \langle\delta\rangle=\frac{1}{N}\sum_i \delta_i} \max_i K_{ii}\delta_i \tag{S6}$$

Here we can leverage the decoupling property of our population decline criterion. By assigning the maximum possible relative growth to the microhabitat with the highest inversed dynamic-related centrality $K_{ii}$, we obtain the condition:

$$\max K_{ii} \max \delta_i < 1$$

For a numerical result of the phase boundary, we can numerically compute $\max K_{ii}$ directly and construct a phase diagram, as illustrated in the main text, to visualize the transition from decline to other regimes.

#### Sufficient condition of robust population decline by linear interpolation upper bound

For robust population decline regardless of the spatial drug heterogeneity, we require

$$\max K_{ii} \max \delta_i < 1$$

where $\max \delta_i = 1 + \frac{\min(g_0, N\langle g\rangle-(N-1)d_0)}{|d_0|}$ is the maximum possible relative growth deviation. Recall $K_{ii} = \sum_{k=1}^N \frac{V_{ik}\left(V^{-1}\right)_{ki}}{1+\frac{\beta}{|d_0|}\omega_k}$. We can treat $V_{ik}\left(V^{-1}\right)_{ki}$ as weights, denoted by $p_k = V_{ik}\left(V^{-1}\right)_{ki}$, for which we have $\sum_{k=1}^N p_k = 1$. The sum of weighting factors $\sum_k V_{ik}\left(V^{-1}\right)_{ki} = 1$ since $VV^{-1} = I$ - it's the ith diagonal entry of identity matrix. Thus $K_{ii} = \sum_{k=1}^N \frac{V_{ik}\left(V^{-1}\right)_{ki}}{1+\frac{\beta}{|d_0|}\omega_k} = \sum_{k=1}^N p_k f(w_k)$, where $f(x) = \frac{1}{1+\frac{\beta}{|d_0|}x}$ and it's a convex function by observation. In the context of weighted averages for convex functions, we have an inequality expressed as an linear interpolation between the minimum and maximum values of the function. Specifically:

$$\sum_{k=1}^N p_k f(x_k) \leq \frac{f(M)-f(m)}{M-m}\left(\sum_{k=1}^N p_k x_k - m\right) + f(m) \tag{S7}$$

where $m$ is the minimum value and $M$ is the largest value. Apply it to our inversed centrality $C_i^{-1} = K_{ii}$, we have

$$\begin{aligned} K_{ii} \leq& \frac{f(M)-f(m)}{M-m}\left(\sum_{k=1}^N p_k x_k - m\right) + f(m) \\ =& 1 - \frac{1-f(\omega_{max})}{\omega_{max}} k(i) \end{aligned}$$

$k(i)$ is the degree of the ith microhabitat; for directed graph, it's the out degree from the ith microhabitat. To guarantee the robust population decline, here we require $\max K_{ii} \max \delta_i < 1$. So if $(1-\frac{1-f(\omega_{max})}{\omega_{max}} k_{min}) \max \delta_i < 1$, the population decline is guaranteed and this inequality serves as a sufficient condition. By doing the re-arrangement, we have

$$\beta > \frac{|d_0|}{k_{min}(1+\frac{1}{\delta-1})-\omega_{max}} \tag{S8}$$

where $\delta = \max_i \delta_i = 1 + \frac{\min(g_0, N\langle g\rangle-(N-1)d_0)}{|d_0|}$.

For equality to hold in the linear interpolation inequality, the following conditions must be satisfied: all the arguments $x_k$ are either equal to the minimum value $m$ or the maximum value $M$. This means that the points $x_k$ must lie at the extremes of the interval $[m, M]$. In other words, there must be a partition of the weights such that:

$$x_k = m \quad \text{for some } k \quad \text{or} \quad x_k = M \quad \text{for others}$$

In the language of graph, this requires either $w_k = 0$ or $w_k = w_{max}$. The only graph that satisfies such restrictions is the complete graph $K_N$. So, for complete graphs, our sufficient condition becomes both sufficient and necessary, thus exact. We can also know that graphs with a high multiplicity of eigenvalues at one or both extremes (like cycle graphs, star graphs, or regular graphs with highly symmetric spectra) are good candidates for being close to the upper bound and making the condition near exact.

#### Necessary condition of robust population decline by Jensen's inequality

Similarly, we can also derive a necessary condition for population decline by Jensen's inequality. Jensen's inequality states that for a convex function $f$ :

$$f\left(\sum_{k=1}^{N} p_k x_k\right) \leq \sum_{k=1}^{N} p_k f\left(x_k\right) \tag{S9}$$

if $f$ is convex and $p_k \geq 0$ with $\sum_{k=1}^{N} p_k = 1$. Applying Jesen's inequality to $K_{ii}$:

$$K_{ii} = \sum_{k=1}^{N} p_k f\left(\frac{\beta}{|d_0|}\omega_k\right) \geq f\left(\sum_{k=1}^{N} p_k \frac{\beta}{|d_0|}\omega_k\right)$$

Using the convexity property, we have:

$$K_{ii} \geq \frac{1}{1+\sum_{k=1}^{N} p_k \frac{\beta}{|d_0|}\omega_k}$$

To proceed further, let's denote: $\langle\omega\rangle_i = \sum_{k=1}^{N} p_k \omega_k$, which represents a weighted average of the $\omega_k$ at the ith microhabitat. We can also interpret $\langle\omega\rangle_i$

in another way:

$$\langle\omega\rangle_i = \sum_{k=1}^{N} V_{ik}\omega_k \left(V^{-1}\right)_{ki} = L_{ii}$$

is actually the diagonal entry of the product $V\Lambda V^{-1}$, specifically for row and column $i$. Therefore, $\bar{\omega}$ represents the value of $L_{ii}$, the $i$-th diagonal entry of the Laplacian matrix which is also the degree of ith microhabitat. Then:

$$K_{ii} \geq \frac{1}{1+\frac{\beta}{\alpha}k(i)}$$

For the maximum of $K_{ii}$ :

$$\max K_{ii} \geq \frac{1}{1+\frac{\beta}{|d_0|}k_{min}}$$

Thus we derive a necessary bound for $C_i$ :

$$\frac{1}{1+\frac{\beta}{|d_)|}k_{min}} \max \delta_i \leq \max C_i \max \delta_i < 1$$

So

$$\beta > \frac{(\delta-1)|d_0|}{k_{min}} \tag{S10}$$

where $\delta = \max_i \delta_i = 1 + \frac{\min(g_0, N\langle g\rangle-(N-1)d_0)}{|d_0|}$.

For Jensen's inequality to hold with equality in this specific case, one of the following conditions must be true: 1. all arguments are Same - all $\frac{\beta}{|d_0|}\omega_k$ are equal for all $k$. However, this is not possible since for any graph with a single component, the minimum eigenvalue of Laplacian matrix is 0. And the rest eigenvalues are non-zero. 2. the function $f(x) = \frac{1}{1+x}$ must be linear over the interval of the inputs. However, in this scenario it's impossible. 2 robust decline phase digrams with both sufficient and necessary conditions are shown, with cycle graph and star graph as examples(See Figure S2, Figure S3).

### Comparison between dynamic-related centrality $C$, closeness centrality, and other centrality measures

Centrality measures are fundamental tools for quantifying node importance in a network. Our proposed dynamic-related centrality $C_i = \frac{1}{K_{ii}}$ shares mathematical similarities with the forest closeness centrality $C_i^F = \frac{N}{K_{ii}+\mathrm{tr}(K)-2}$(See Maintext, Discussion section), as both are derived from diffusion-based dynamics and capture global structure information through spectral properties of the Laplacian matrix.

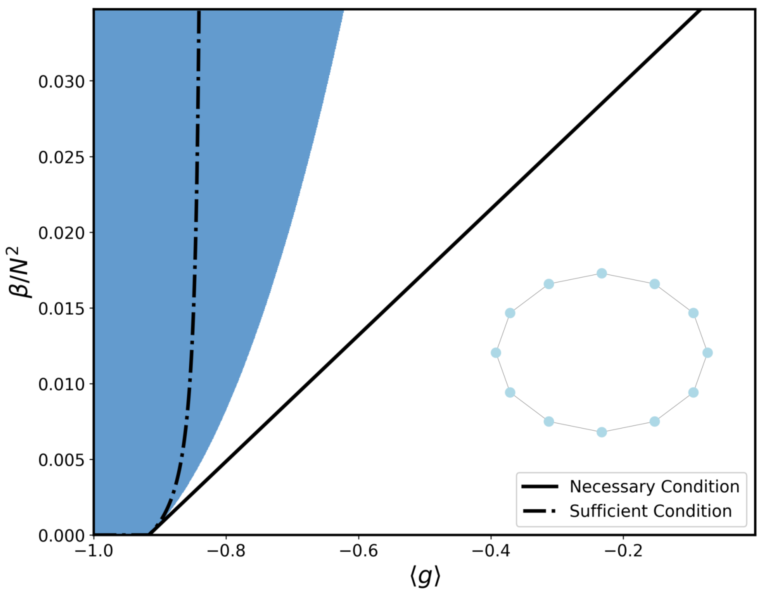


Figure S2: **Phase diagram of migration rate $\beta$ and spatially averaged growth rate $\langle g \rangle$ on a cycle graph as the microhabitat structure.** The dashed line and solid line are sufficient and necessary conditions for robust population decline(blue phase). $N = 12, d_0 = -1, g_0 = 22$.

To compare our proposed dynamic-related centrality $C_i$ with traditional network centralities, we visualize different centrality metrics in Figure S4, examine their correlations in Figure S5, and analyze their pairwise relationships in Figure S6.

In Figure S4, dynamic-related centrality $C_i$ exhibits a strongly similar pattern to Degree Centrality, Katz Centrality, and PageRank Centrality (correlation score $r = 0.99$ shown in Figure S5), followed closely by Eigenvector Centrality (0.95). This is consistent with our earlier analytical approximation, which suggested that $C_i$ scales with degree under certain assumptions. These results also suggest that, in networks with complex microhabitat structures, targeting nodes with extreme degree values may serve as a visually interpretable and suboptimal—but practical-clearance strategy.

Interestingly, $C_i$ and Forest Closeness Centrality $C_i^F$, despite both being based on Laplacian regularization, only show a moderate linear correlation (0.90). This difference is more pronounced when compared to the relatively low correlation between $C_i$ and classical Closeness Centrality (0.83), and a slightly higher one with Betweenness Centrality (0.89). These suggest that although $C_i$ shares structural similarities with other Laplacian-based or path-based measures, it captures distinct dynamics-driven features.

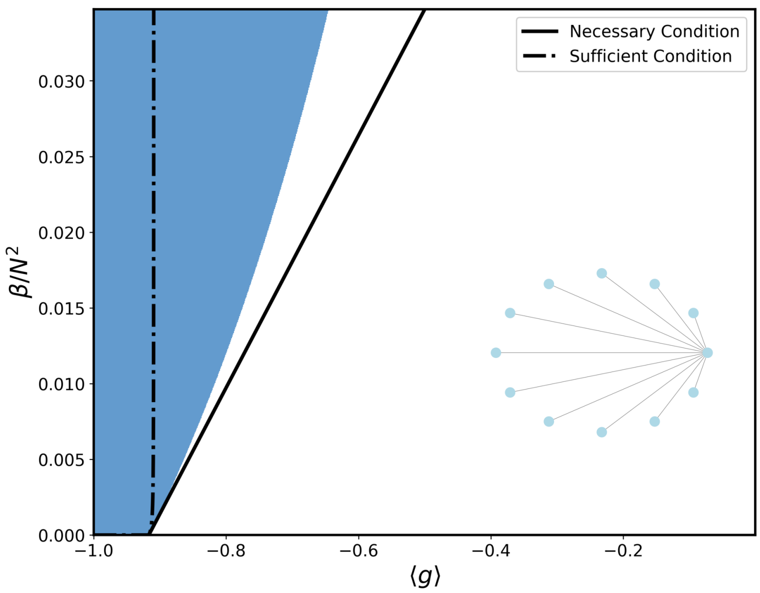


Figure S3: **Phase diagram of migration rate $\beta$ and spatially averaged growth rate $\langle g \rangle$ on a star graph as the microhabitat structure.** The dashed line and solid line are sufficient and necessary conditions for robust population decline(blue phase). $N = 12, d_0 = -1, g_0 = 22$.

Similar trends are evident in the scatter plot matrix (Figure S6). Notably, while $C_i$ and Forest Closeness Centrality display only moderate linear correlation, their scatter plot reveals a visually tight nonlinear relationship, indicating that they may produce nearly identical node rankings despite a lower Pearson coefficient. This underscores a key limitation of linear correlation: it fails to capture monotonic or nonlinear alignments.

Our findings suggest that dynamic-related centrality $C$ provides a complementary perspective distinct from classical centrality measures, making it particularly relevant in modeling network stability and dynamical processes such as population decline, disease spreading, and diffusion dynamics. This property also aligns with DomiRank[23], a centrality framework that models node importance through dynamical processes, though unlike our approach, it assumes homogeneous node properties rather than incorporating heterogeneous effects as considered in $C$.

#### Supplementary derivations

**Diagonal matrix** Let's assume 3 different matrices $A, B, C$, where $A, C$ are diagonal matrices, for $ABC$, we have

$$
\begin{aligned}
(ABC)_{ij} &= \sum_k \sum_l a_{ik} b_{kl} c_{lj} \\
&= \sum_k \sum_l a_{ik} \delta_{ik} b_{kl} c_{lj} \delta_{lj} \\
&= a_{ii} b_{ij} c_{jj}
\end{aligned}
$$

**Sufficient criterion of population decline by Gershgorin's Circle Theorem** Here let's restate the Theorem first: Let $A \in \mathbb{R}^{n \times n}$. Then all eigenvalues of $A$ lie within the union of the Gershgorin disks:

$$
\lambda \in \bigcup_{i=1}^{n} \left\{ z \in \mathbb{C} \mid |z - A_{ii}| \leq \sum_{j \neq i} |A_{ij}| \right\}
$$

So if every disk lies strictly inside the unit disk, then

$$
\lambda_{\max}(A) < 1 \quad \Rightarrow \quad A \prec I
$$

Let's compute $A_{ij} = \sqrt{\delta_i \delta_j} K_{ij}$. Then the $i$-th Gershgorin disk for $A$ is:

$$
\left\{ z \in \mathbb{C} \mid |z - \delta_i K_{ii}| \leq \sum_{j \neq i} \sqrt{\delta_i \delta_j} |K_{ij}| \right\}
$$

or more compactly:

$$
\max_i \left( \delta_i K_{ii} + \sum_{j \neq i} \sqrt{\delta_i \delta_j} K_{ij} \right) < 1 \quad \Rightarrow \quad \lambda_{\max}(A) < 1 \Rightarrow \text{ decline}
$$

This is a strictly sufficient condition: if it's satisfied, decline is guaranteed.

**Centrality-based lower bound derivation by Rayleigh quotient** The dominant eigenvalue can be rewritten as the Rayleigh quotient

$$
\max \lambda(A) = \max_x \frac{x^\top A x}{x^\top x}
$$

where $x^\top A x = \sum_{i,j} x_i x_j A_{ij}$. By the property of Rayleigh quotient, we have

$$
\max \lambda(A) \geq \frac{x^\top A x}{x^\top x}
$$

for an arbitrary test vector $x$. When $x = v_{\max}$, the eigenvector corresponding to the largest eigenvalue, the equality holds. If we choose $x = \max\{e_1, e_2, \cdots, e_N\}$, then we get

$$\max \lambda(A) \geq \max_i \frac{e_i^\top A e_i}{e_i^\top e_i} = \max_i K_{ii}\delta_i, i = 1, 2, ..., N$$

This is exactly the centrality-based lower bound. If we choose $x = \frac{1}{\sqrt{N}}\mathbf{1}$, then we get a lower bound

$$\max \lambda(A) \geq \sum_{i,j} x_i x_j A_{ij}$$

This can be treated as a all-sum version of the upper bound of $\lambda_{\max}(A)$ derived before.

As shown in the derivation of the upper bound of $\lambda_{\max}(A)$, we can also improve the centrality-based lower bound (or sufficient condition for population growth) by iteratively improving the test vector $x$ from an optimization perspective, with methods such as power iteration, or Lanczos methods. Here we choose power iteration as an example. To estimate $\lambda_{\max}(A)$, pick any vector $x_1 \neq 0$ and repeatedly apply

$$x_{k+1} := \frac{Ax_k}{\|Ax_k\|}$$

This converges to the dominant eigenvector $v_{\max}$ of $A$. Then:

$$\lambda_{\max}(A) = \lim_{k\to\infty} \frac{x_k^\top A x_k}{x_k^\top x_k}$$

Again, for the first step, if we choose the test vector as $x_1 = \max\{e_1, e_2, \cdots, e_N\}$, we can get $\lambda_{\max}(A) \geq \max K_{ii}\delta_i$. For $k \geq 2$, by applying the iteration function, we can get a more and more tight loewr bound until it becomes exact.

**Why always $K_{ij} > 0$** Because we have $L$ is positive semidefinite, $I + \alpha L$ is strictly positive definite → its inverse exists. The inverse of a strictly diagonally dominant matrix like $I + \alpha L$ is entrywise positive. So:

$$K = (I + \alpha L)^{-1} \quad \Rightarrow \quad K_{ij} > 0 \quad \forall i, j$$

Even for nodes with no direct path in the original graph - there's indirect interaction through the Laplacian spectrum.

**Diagonal dominance of $K$** First let's point out that $(I + \alpha L)$ is strictly diagonally dominant. For a Laplacian matrix $L$, because $(I+\alpha L)_{ii} = 1+\alpha \deg(i)$ and sum of off-diagonal magnitudes is $\alpha \deg(i)$, we have

$$(I + \alpha L)_{ii} = 1 + \alpha \deg(i) > \alpha \deg(i) = \sum_{j\neq i} |(I + \alpha L)_{ij}|$$

By applying the following theorem we can find out that, the inverse of a strictly row diagonally dominant matrix has a form of diagonal dominance, namely that the largest element in each column is on the diagonal. This theorem and its proof can be found out in Professor Nick Higham's blog (nhigham.com). Since the theorem is true, so $K$ has dominant diagonal terms and $K_{ii} > K_{ij}$.

**Theorem 2.** *If $A \in \mathbb{C}^{n\times n}$ is strictly diagonally dominant by rows then $B = A^{-1}$ satisfies $|b_{ij}| < |b_{jj}|$ for all $i \neq j$.*

**Proof** For $i \neq j$ we have $\sum_{k=1}^{n} a_{ik} b_{kj} = 0$. Let $\beta_j = \max_k |b_{kj}|$. Taking absolute values in $a_{ii} b_{ij} = -\sum_{k\neq i} a_{ik} b_{kj}$ gives

$$|a_{ii}|\,|b_{ij}| \leq \beta_j \sum_{k\neq i} |a_{ik}| < \beta_j\,|a_{ii}|$$

or $|b_{ij}| < \beta_j$, since $a_{ii} \neq 0$. This inequality holds for all $i \neq j$, so we must have $\beta_j = |b_{jj}|$ , which gives the result.

**Functions of Laplacian matrix $L$ preserve symmetry** Now consider a function of $L$ - such as $f(L) = (I + \alpha L)^{-1}$ or any other analytic function like $\exp(-\beta L)$, etc. We have such a fact from linear algebra: for permutation matrix $P$, if $P^\top LP = L$, then for any matrix function $f$, $P^\top f(L)P = f(L)$, (i.e., $f(L)$ commutes with the graph symmetry). So for our case:

$$K = (I + \alpha L)^{-1} \quad \Rightarrow \quad P^\top KP = K$$

That means $K$ preserves all the same structural symmetries as $L$.

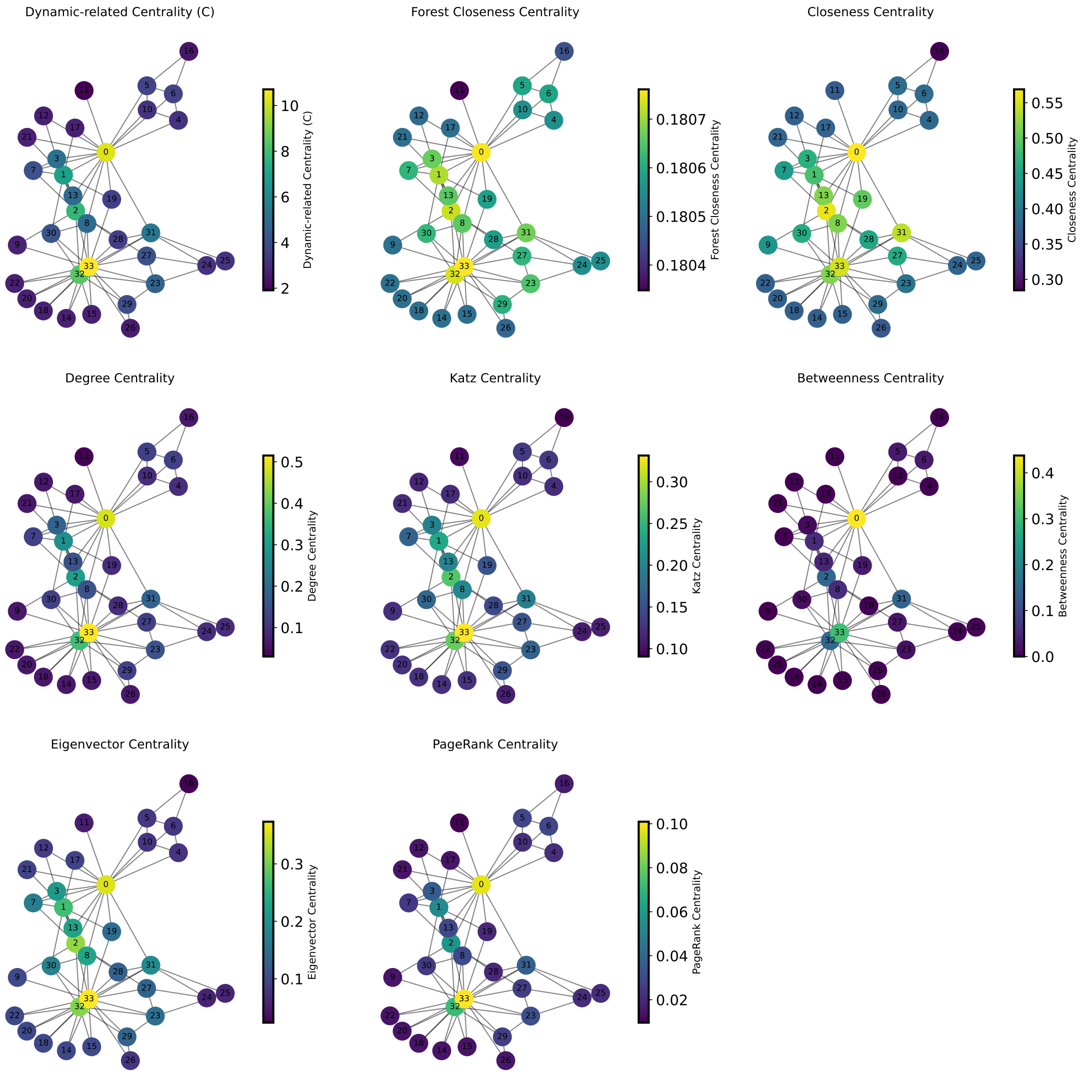


Figure S4: **Comparison between dynamic-related centrality $C$, forest closeness centrality, and other centrality measures in a network(Karate Club network).** Each subplot represents a different centrality metric, illustrating node importance based on different criteria. Nodes are colored according to their centrality values, with yellow indicating higher centrality and purple representing lower centrality. The metrics include (top row) dynamic-related centrality (C), forest closeness centrality, and closeness centrality; (middle row) degree centrality, Katz centrality, and betweenness centrality; (bottom row) eigenvector centrality and PageRank centrality. Notably, the dynamic-related centrality (C), degree centrality, and PageRank centrality exhibit similar patterns. For parameters, $\gamma = 1$ for dynamic-related centrality and forest closeness centrality, $\alpha_{Katz} = 0.1$.

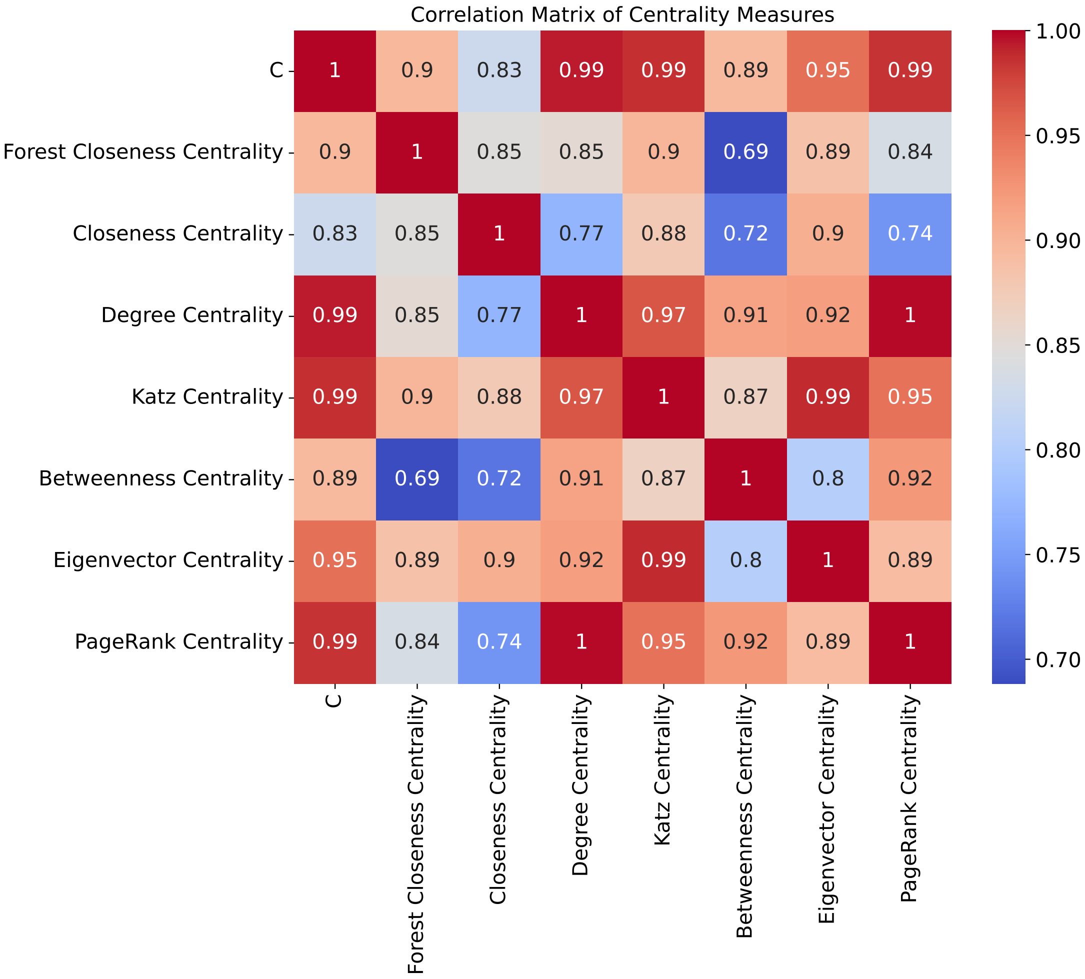


Figure S5: **Correlation matrix of different centrality measures in the network(Karate Club network).** The heatmap shows the Pearson correlation coefficients between different centrality metrics, where red indicates strong positive correlations and blue indicates strong negative correlations. Notably, dynamic-related centrality (C), degree centrality, Katz centrality, and PageRank centrality are almost perfectly correlated ($r = 0.99$), suggesting they capture the same structural properties. The dynamic-related centrality only has a correlation coefficient at $r = 0.90, 0.83$ with forest closeness centrality and closeness centrality.

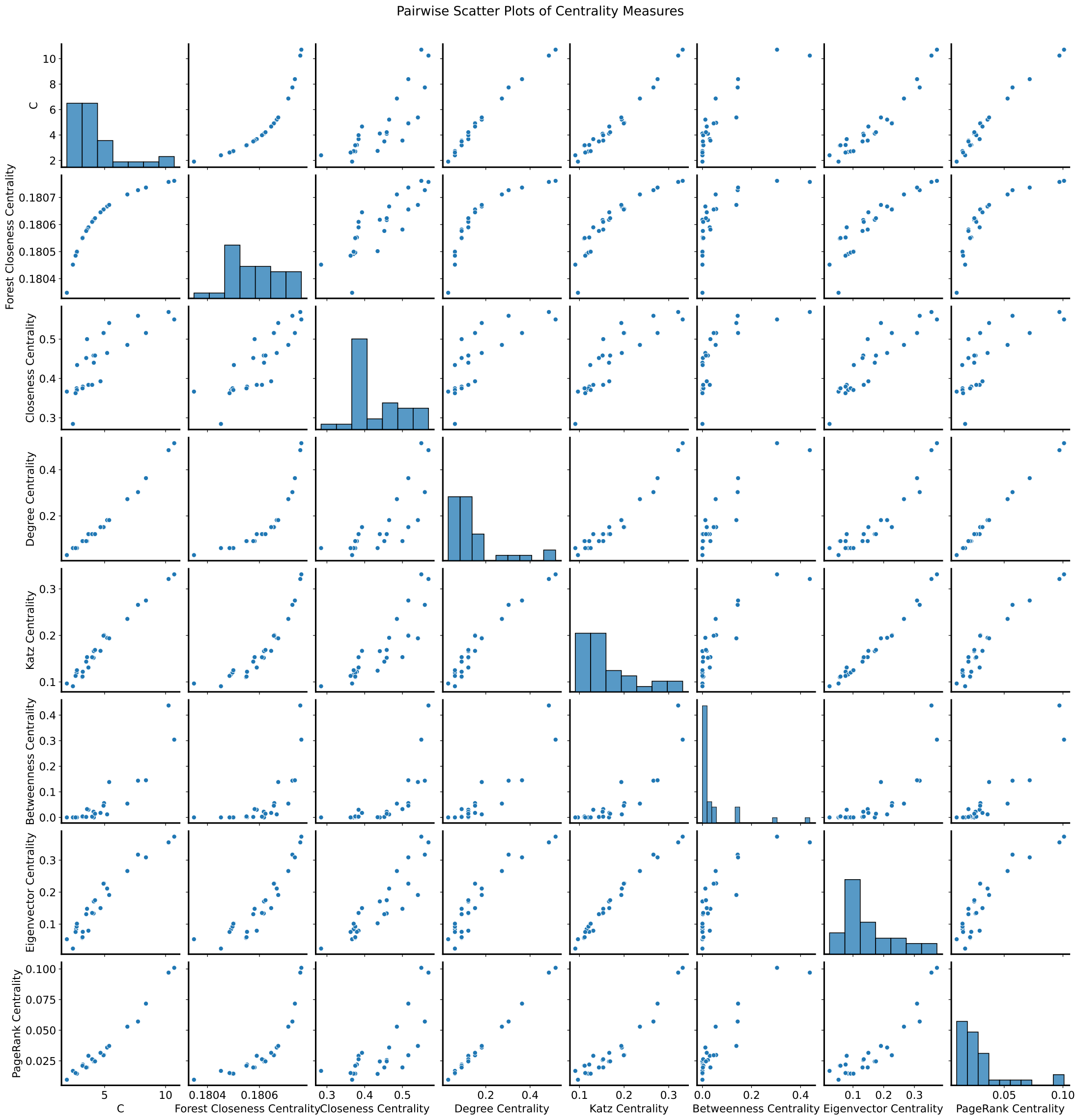


Figure S6: **Pairwise scatter plots of centrality measures in the network.** Each subplot compares two centrality metrics, with histograms along the diagonal showing the distribution of each measure. Strong positive correlations are observed between dynamic-related centrality (C), degree centrality, Katz centrality, and PageRank centrality.
